# Tween-20 induces the structural remodelling of single lipid vesicles

**DOI:** 10.1101/2022.03.01.482482

**Authors:** Lara Dresser, Sarah P. Graham, Lisa M. Miller, Charley Schaefer, Donato Conteduca, Steven Johnson, Mark C. Leake, Steven D. Quinn

## Abstract

The interaction of Tween-20 with lipid membranes is crucial for a number of biotechnological applications including viral inactivation and membrane protein extraction, but the underlying mechanistic details have remained elusive. Evidence from ensemble assays supports a global model of Tween-20 induced membrane disruption that broadly encompasses association of the surfactant with the membrane surface, membrane fragmentation and the release of mixed micelles to solution, but whether this process involves intermediate and dynamic transitions between regimes is an open question. In search of the mechanistic origins of membrane disruption, increasing focus is put on identifying Tween-20 interactions with highly controllable model membranes. In light of this, and to unveil quantitative mechanistic details, we employed highly interdisciplinary biophysical approaches, including quartz-crystal microbalance with dissipation monitoring, steady-state and time-resolved fluorescence and FRET spectroscopy, dynamic light scattering, fluorescence correlation spectroscopy, wide-field single-vesicle imaging and scanning electron microscopy, to interrogate the interactions between Tween-20 and both freely-diffusing and surface-immobilized model-membrane vesicles. Using ultrasensitive sensing approaches, we discovered that Tween-20 leads to a stepwise and phase-dependent structural remodelling of sub-micron sized vesicles that includes permeabilization and swelling, even at detergent concentrations below the critical micellar concentration. These insights into the structural perturbation of lipid vesicles upon Tween-20 interaction highlight the impact on vesicle conformation prior to complete solubilization, and the tools presented may have general relevance for probing the interaction between lipid vesicles and a wide variety of disruptive agents.

## Introduction

Tween-20 is a non-ionic detergent widely used as a solubilizing agent of membrane proteins(1, 2) and for the inactivation of enveloped viruses(3, 4). Previous findings have also suggested that the detergent substantially enhances drug permeability through cell membranes(5, 6), and regulates the structure and diffusion properties of transmembrane proteins(7). However, despite such important roles across a number of applications, the mechanistic details of the Tween-20-membrane interaction remain largely ill-defined.

While the detergent is often used empirically as a general membrane solubilizer(2), experiments involving giant unilamellar vesicles (GUVs) have been employed as a means to explore the underlying details. The biophysical properties of GUVs, which mimic the cell membrane, are highly controllable synthetic models that provide opportunities for interrogating detergent-membrane interactions in the absence of extraneous biochemical processes(8). For instance, high-intensity dark-field microscopy has enabled structural changes in GUVs ∼5 μm in size to be observed in response to Tween-20, revealing morphological events, such as stepwise shrinking, vigorous intermittent fluctuations, and catastrophic bursting that are regulated by the lipid composition(9). Similarly, the observation of ∼10 μm sized vesicles revealed that at low detergent concentrations an increase in the membrane surface area prior to pore-forming associated shrinkage occurs(10). More broadly, GUVs undergoing Tween-20 induced solubilization also display the transient and cyclic opening of pores and the rate of solubilization is enhanced with increased surfactant concentration(11). When even larger vesicles (> 10 μm) were placed in a Tween-20 concentration gradient spanning 0 – 0.6 mM, the latter representing ∼10x the reported critical micellar concentration (cmc), the pore lifetime was found to be of the order of minutes(12). In the same study, optical microscopy also revealed that the opening of transient pores facilitated vesicle fusion, though whether oscillatory pore motion, which has been observed for other non-ionic detergents(13), plays a role in this process remains unclear. Due to its ability to regulate membrane elasticity, Tween-20 has found utility in the generation of highly elastic vesicles that facilitate drug transport across mouse and human skin(14) and in niosomes, where non-ionic surfactants such as Tween-20 are the main ingredient, the transport of encapsulated drugs across the intestinal epithelial barrier is enhanced(15, 16). More recently, dynamic light scattering (DLS) and turbidity approaches have been employed to indicate the bending rigidity of vesicle bilayers decreases quasi-exponentially with increasing concentration of the longer chain surfactant Tween-80(17).

On the basis of these experimental results, and others which report substantial alteration of membrane conformation in response to non-ionic detergents more generally(18-20), a global three-step model has been proposed for the mode of Tween-20 induced membrane solubilization. Here, the detergent monomers saturate the membrane in step 1, leading to the formation of mixed detergent-lipid micelles and fragmentation of the membrane in step 2, and release of mixed detergent-lipid micelles to solution in step 3(21). A more quantitative extension of this model involves the formation of transient polar defects and micropores on the in-tact membrane prior to complete solubilization, where such events are dependent on the lipid composition, phase and detergent concentration(22). However, the use of GUVs as model systems only represents one end of the membrane curvature space (23), and the use of conventional optical imaging approaches only allows for macroscopic changes in vesicle shape and packing density of GUVs typically > 5μm to be inferred. Consequently, such experiments provide little detail on the molecular level(24). Given highly curved vesicles, which are an order-of-magnitude smaller than GUVs and not easily visualized by diffraction-limited microscopy techniques, and yet are commonly used in a range of bionanotechnological applications and have important implications in the context of biological trafficking(25, 26), it is important to explore the molecular details of the Tween-20 interaction at the opposite end of the membrane curvature space.

Recent developments in structural and bioanalytical methods have brought the understanding of the molecular mechanisms of detergent-induced membrane disruption forward. For instance, single-vesicle Förster resonance energy transfer (FRET) imaging applied to sub-micron sized vesicles revealed that the detergent Triton-X 100 induces dynamic transitions between regimes in the three step model(23). Similarly, vesicle swelling induced by the ionic detergent sodium dodecyl sulfate (SDS) was also observed by the combined use of FRET, atomic force microscopy and quartz crystal microbalance with dissipation (QCM-D) monitoring(24). Studies using a combination of light scattering, fluorescence correlation spectroscopy (FCS), cryo-electron microscopy and coarse-grained molecular dynamic simulations have also revealed dynamic phase transitions and remodelling during the detergent-membrane interplay(18, 27-31), suggesting the three step model may also involve a number of additional transient, dynamic and interlinked events.

Inspired by these insights, we adopted a highly interdisciplinary approach to interrogate the Tween-20 membrane interaction, focusing specifically on the interaction between Tween-20 and sub-micron sized lipid vesicles of ∼200 nm diameter. While previous single-vesicle studies on GUVs utilized fluorescence alone, we employed a range of tools, including QCM-D to explore mass and viscoelasticity changes, ensemble and time-resolved FRET spectroscopy to assess lipid partitioning, DLS and FCS to probe the hydrodynamic diameters of freely-diffusing vesicles and wide-field total internal reflection fluorescence microscopy and scanning electron microscopy imaging to capture the response from single immobilized vesicles. An important aspect of this work is the use of ensemble and single-vesicle FRET spectroscopy, which is capable of reporting the distance between donor and acceptor probes embedded within the membrane with ∼1 nm spatial resolution. We previously used these techniques to quantify the solubilization dynamics of large unilamellar vesicles in response to TX-100 and SDS, and reveal kinetically asynchronized reductions in FRET efficiency (reflecting vesicle swelling) and reduction in total fluorescence intensity (reflecting lysis)(23, 24). Here, we used these techniques to explore how Tween-20 affects the structural integrity of vesicles composed of POPC (1-palmitoyl-2-oleoyl-glycero-3-phosphocholine) and we extract information about the membrane composition and interaction by applying a mass-action model to variations in the FRET efficiency(32). To further characterize the interaction, we also implemented an ultrasensitive analytical approach to quantify the extent of membrane disruption by Tween-20 whereby vesicles filled with the fluorescent molecule Cal-520, whose fluorescence increases upon binding Ca^2+^ ions, report quantitatively on Ca^2+^ entry into vesicles as a consequence of permeabilization(33).

Our discovery of the structural remodelling of both freely-diffusing and surface-immobilized vesicles in response to Tween-20, even at concentrations below the cmc, contributes new clues to the underlying solubilization mechanism. We also expect the presented tools may have far-reaching applications in elucidating the underlying membrane damage mechanisms associated with a wide-variety of membrane disruptive molecules.

## Results

### Tween-20 alters the viscoelasticity and mass of surface-tethered vesicles

QCM-D was first used to examine time- and concentration-dependent changes in the mass and viscoelasticity induced by Tween-20 binding to surface-immobilized vesicles. The vesicles were modified with 1 mol % of biotinylated lipids and were tethered via NeutrAvidin to a bovine serum albumin (BSA) coated SiO_2_ sensor containing 2 mol % biotinylated-BSA (BSA-Bi), as demonstrated by real-time changes in the resonance frequency shift (ΔF) and energy dissipation (ΔD) which reflect the mass and viscoelasticity of the surface, respectively (**Figure 1A**). After the immobilization procedure, the sensor surface was rinsed with buffer (50 mM Tris, pH 8) to remove unbound vesicles. Upon addition of 0.02 mM Tween-20 to the immobilized vesicles (t = 105 mins), we first observed a ∼10 Hz reduction in ΔF reflecting an increase in the sensor mass, concurrent with a 20% increase ΔD, representing an increase in viscoelasticity (**Figure 1A**). Control experiments performed simultaneously using sensors coated in BSA, biotinylated-BSA and NeutrAvidin but lacking vesicles displayed similar ΔF and ΔD responses when flushed with Tween-20, which we attributed to a combination of the change in buffer and an increase in mass caused by Tween-20 non-specifically binding to the surface (**Figure S1**). However, when the control surface was washed with buffer (50 mM Tris, pH 8), a recovery in ΔF and ΔD to similar levels prior to the introduction of Tween-20 was observed, indicating the detergent does not lead to the release of BSA-Biotin or NeutrAvidin, and the non-specific attachment of Tween-20 is reversible. The rate of change across both signals was ∼ 3x faster when vesicles were present, pointing towards a more rapid mass and viscoelastic gain. The subsequent and dramatic increase in ΔF at t ∼ 110 mins, as seen on the vesicle-coated surface (**Figure 1A**), and the anti-correlated decrease in ΔD, can then be explained by the removal of lipid material and surfactant from the substrate with a rate constant of 0.25 ± 0.01 Hz min^-1^. Similar behaviour was observed when 0.04 mM and 0.06 mM Tween-20 was flushed across vesicle-coated surfaces (**Figures 1B and 1C**), indicating similar degrees of mass gain and comparable solubilization rates (0.27± 0.01 Hz min^-1^). Under the latter conditions we observed a further reduction in ΔF and corresponding increase in ΔD after the major mass loss event that we assigned to the adsorption of mixed detergent-lipid micelles on the surface. The Tween-20 interaction with the vesicle-layer was also visualized by plotting changes in ΔF against the corresponding changes in ΔD across the timescale of the interaction, with the interaction defined as complete when the plateau region of the QCM-D response was ΔF/Δt < 2 Hz / 10 min (**Figure 1D**). Higher negative frequency shifts were assigned to an increase in adsorbed mass, whereas higher dissipation values reflect increasing viscoelasticity. Under each condition tested, the ΔF versus ΔD responses displayed an initial turning point, reflecting a mass gain and increase in viscoelasticity, followed by substantial mass loss to the solution, which we assigned to lysis. Taken together, the QCM-D data indicates that the initial deposition of Tween-20 onto the surface triggers a substantial structural remodelling event that precedes the loss of material to solution. Indeed, the temporal evolution of the surface upon the addition of Tween-20 under all conditions tested agrees with this assessment, with structural variations found to occur on broadly similar timescales across the concentrations tested. A surprising outcome of this analysis is that the initial structural rearrangement occurred at concentrations of detergent below the cmc (∼0.06 mM), pointing towards an interaction between detergent monomers and vesicles that leads to substantial conformational changes prior to lipid release.

**Figure 1.**
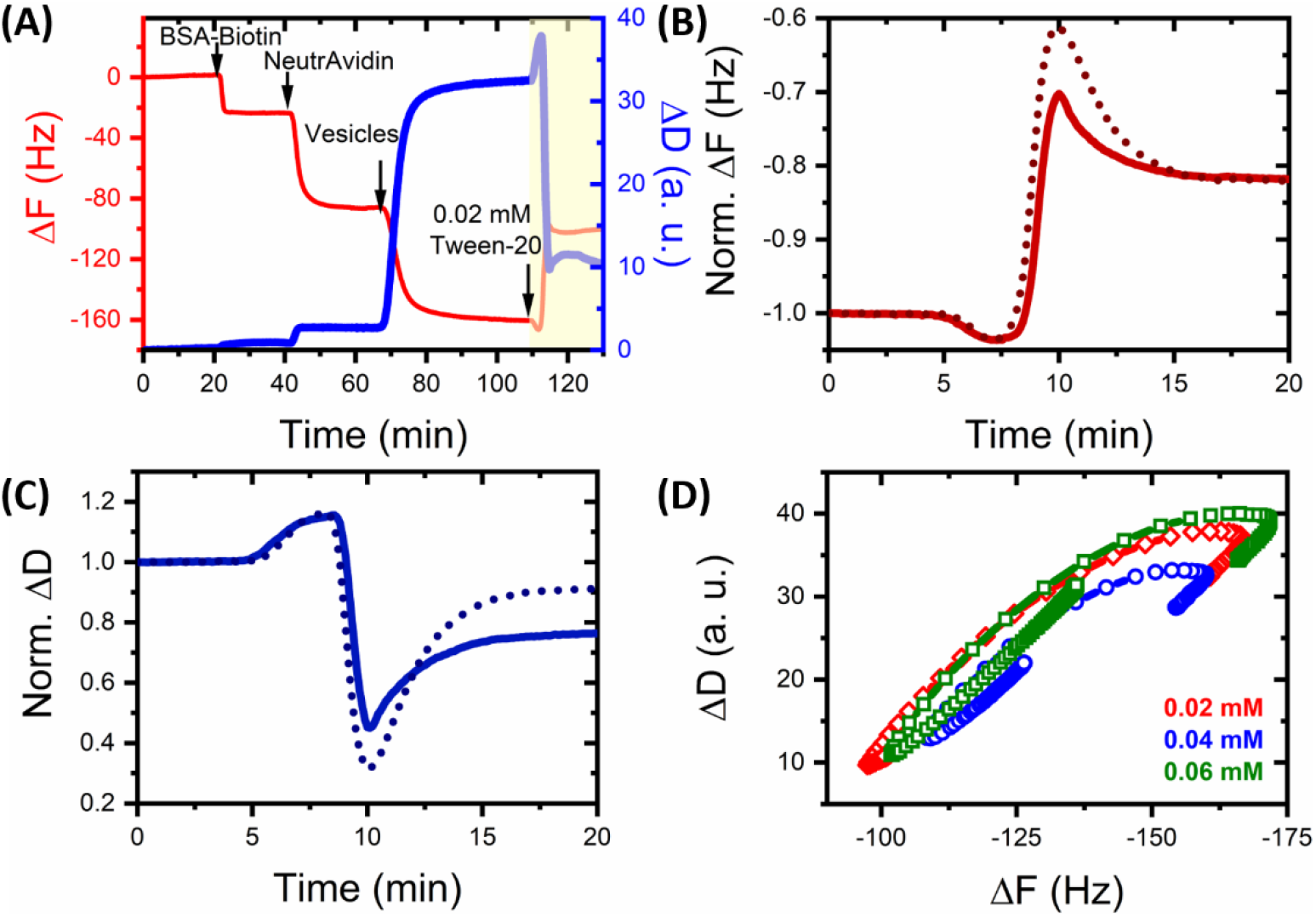
QCM-D of Tween-20 interactions with surface-tethered lipid vesicles. (A) Time evolution of ΔF (red) and ΔD (blue) upon the addition of POPC vesicles to a BSA-biotin and NeutrAvidin-coated sensor surface. Addition of Tween-20 at 0.02 mM (yellow shaded area) was performed after washing the vesicle-saturated surface with 50 mM Tris buffer (pH 8). Normalized variations in (B) ΔF and (C) ΔD observed in response to 0.04 mM (solid lines) and 0.06 mM (dashed lines) Tween-20 injected at t ∼ 5 min. (D) Frequency versus dissipation observed during the interaction between surface immobilized vesicles and Tween-20 under the conditions tested. Data is shown in all cases for the 7^th^ harmonic.

### Tween-20 Induces the Structural Remodelling of Freely-Diffusing POPC Vesicles

To gain molecular-level insights into the observed structural changes, we next performed steady-state and time-resolved Förster resonance energy transfer (FRET) experiments. Here, we recorded the fluorescence emission spectra and time-resolved fluorescence decays obtained from freely-diffusing lipid vesicles incorporating the lipophilic fluorescent dyes DiI and DiD (**Figure 2A**). As we previously reported, the incorporation of 0.1 mol% DiI (FRET donor) and 0.1 mol% DiD (FRET acceptor) into 200 nm sized lipid vesicles, leads to an average FRET efficiency per vesicle, E, of ∼0.5. Given E is inversely proportional to the sixth power of distance between the probes, structural variations such as swelling or compaction, are reported through changes in E in either direction with a spatial resolution of ∼1 nm. Using this approach, both the emission spectra and time-resolved fluorescence decays were monitored upon addition of Tween-20 above and below the cmc. As shown in **Figure 2B**, the changes in the fluorescence emission spectra before and after addition of Tween-20 are substantial: we observed a two-fold increase in the peak DiI fluorescence emission intensity concurrent with a loss of sensitized DiD emission as Tween-20 was progressively titrated towards 0.3 mM. These observed changes translate to an overall reduction in FRET efficiency and increase of ∼58 % in the mean donor-acceptor separation distance across the titration (**Figure 2B**). The solubilization of the membrane by Tween-20 becomes more prevalent at higher temperatures, as revealed by the shift of the half maximal concentration of the FRET efficiency to lower concentrations at increasing temperatures. Following our previous approach, we quantified this by curve-fitting the Hill model(23) to the data, which yielded a half maximal concentration of 0.11 ± 0.01 mM at 4 °C, ∼0.04-0.06 mM at 21 °C (similar to the reported cmc (34)) and 0.03 ± 0.01 mM at 37 °C. The Hill coefficient was 4.1 ± 0.9 and independent of the temperature. From these descriptive features we extracted physical information about the surfactant-lipid interactions by adopting a mass-action model (**Supplementary Text 1, Figure S2**). The mass-action model predicts that the sharpness of the FRET efficiency curve (which the Hill model describes using the Hill coefficient) is controlled by the excess of free surfactants with respect to the lipids near the solubilization concentration. The model further describes the temperature-controlled shift of the solubilization concentration in terms of a surfactant-to-lipid-membrane binding energy of 31 ± 3 kJ/mol, which is modified for the increasing surfactant-to-lipid ratio in the membrane at increasing concentrations using a Flory-Huggins free energy of mixing. From our curve-fits we extracted a near-athermal Flory-Huggins parameter, χ, of 1.2 ± 0.2 (**Table S1**). As χ=0 indicates ideal miscibility and χ>2 complete incompatibility, the value χ ≈1.2 indicates some limited surfactant-lipid compatibility, which is in line with the idea that the surfactant might structure within the membrane and potentially lead to pore formation. It is also important to emphasize that the reproducibility of the apparent FRET efficiency for in-tact vesicles at 0 mM Tween-20 provides a quantifiable metric to ensure that samples of starting material are identical. We also note that under all conditions tested, a similar final FRET efficiency of ∼0.05 was observed, indicative of a similar final end-state. The overall change in FRET efficiency, consistent with an increase in the average spatial separation of the probes, was assigned to vesicle expansion, micellization or the combination of both in the ensemble.

**Figure 2.**
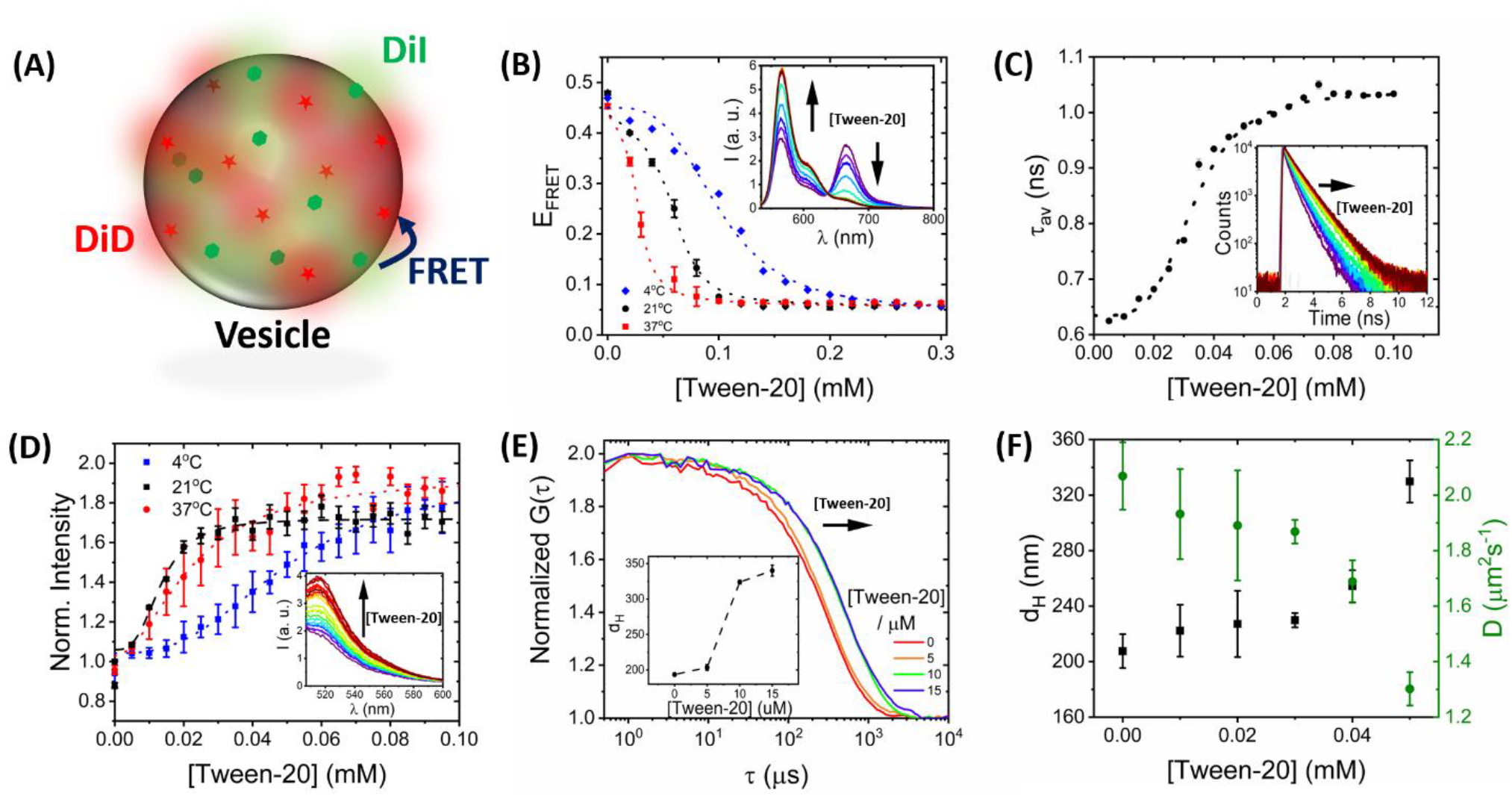
Tween-20 Induces the structural remodelling of freely-diffusing vesicles. (A) Schematic representation of 200 nm sized vesicles composed of 99.8 % POPC, 0.1 % DiI (FRET donor) and 0.1 % DiD (FRET acceptor). (B) Ensemble FRET efficiency of POPC vesicles incorporating DiI and DiD at 4°C (blue), 21°C (black) and 37°C (red) as a function of Tween-20. The dashed lines correspond to mass-action model fits. Inset: representative variation in the fluorescence spectra showing anti-correlative behaviour between the DiI and DiD response. (C) The amplitude weighted average lifetime of DiI, in the presence of DiD, as a function of Tween-20 at 21°C. Inset: the corresponding time-resolved fluorescence decays and instrumental response function (grey). (D) Tween-20 stimulated calcium responses of POPC vesicles encapsulating Cal-520 at 4°C (blue), 21°C (black) and 37°C (red). Inset: the corresponding fluorescence emission spectra of Cal-520 loaded vesicles versus Tween-20. (E) Normalized variation in the DLS correlation curves obtained from DiI labelled vesicles as a function of Tween-20. Inset: variation in the mean vesicle hydrodynamic diameter. (F) Variation in the hydrodynamic diameter (black squares) and diffusion coefficient (green circles) of single vesicles as reported by FCS. Solution conditions: 50 mM Tris, pH 8.0.

To further confirm the presence of an energy transfer mechanism, we next investigated the fluorescence lifetime of DiI in the presence of DiD via time-correlated single photon counting. Here, the amplitude-weighted average lifetime progressively increased as a function of Tween-20, consistent with a progressive enhancement of the donor signal and corresponding decrease in E (**Figure 2C**). Under all conditions, the fluorescence decays were best fitted to a tri-exponential decay model after reconvolution with the instrument response function (**Figure S3**) which as previous studies indicate, likely represents vesicles with coexisting phases in the ensemble, and asymmetry within the membrane(35-38). Our initial studies were performed at 21°C, and in the absence of Tween-20, where the two long-lived environment-sensitive lifetimes were t_1_ = 0.79 ± 0.02 ns (17 %) and t_2_ = 0.34 ± 0.01 ns (16 %), respectively, we recorded an amplitude weighted average lifetime of 0.62 ± 0.01 ns, representative of a quenched DiI state. At 0.1 mM Tween-20, however, the average lifetime increased to 1.03 ± 0.01 ns, and t_1_ and t_2_ increased by > 30 % of their initial values (**Figure 2C**). At 4°C, the initial lifetime at the start of the titration was four-fold longer at 0.76 ± 0.01, likely due to an increase in the fraction of the phase-sensitive components, and a 2-fold increase in average lifetime was observed across the titration (**Figure S4**). At 37°C, a similar trend in the average lifetime was observed, and a half-maximal concentration of 0.02 mM, comparable to those obtained via intensity-based FRET measurements, was observed (**Figure S4**). Taking both sets of measurements together, the data points towards substantial fluorophore separation taking place within the ensemble, though whether this was due to expansion and/or micellization required further investigation. Nevertheless, at 4°C, 21°C and 37°C, the mean spatial separation distance was found to increase by 56 %, 58 % and 52 %, respectively.

We next applied an analytical approach based on the measurement of Ca^2+^ entry into vesicles to further quantify the magnitude of membrane disruption and assess the degree of solution exchange between the vesicle interior and exterior. We employed the use of vesicles loaded with the calcium indicator Cal-520, whose fluorescence emission intensity increases upon binding Ca^2+^. As demonstrated previously in the context of protein-induced membrane disruption(33, 39), Ca^2+^ flux into vesicles as a consequence of membrane permeabilization leads to an increase in the local Ca^2+^ concentration in each vesicle, triggering a concentration-dependant increase in Cal-520 emission. Here, 200 nm-sized POPC vesicles loaded with Cal-520 were prepared as described in the Methods. Non-encapsulated molecules were removed by size exclusion chromatography (**Figure S5**). The loaded vesicles were then incubated in Tween-20 rich solutions and the fluorescence emission spectra were recorded with an excitation wavelength of 493 nm. As the concentration of Tween-20 was progressively increased at 21°C, a two-fold increase in the peak Cal-520 emission intensity at 514 nm was observed (**Figure 2D**). The fluorescence signals before Tween-20 addition were invariant and were found to mono-exponentially increase with a rate constant of 0.10 ± 0.01 s^-1^ after addition of Tween-20 (**Figure S6**). To estimate the mean Ca^2+^ influx into the vesicles, we also added the cation transporting ionophore, ionomycin, enabling the magnitude of Ca^2+^ influx caused by Tween-20 to be unveiled. We observed that the Cal-520 intensity enhancement at 0.06 mM Tween-20 was similar to that observed in the presence of 1 mg/mL ionomycin (**Figure S7**), thus the mean increase in Ca^2+^ influx was found to be ∼95 %. Control experiments performed independently indicated that (i) Ca^2+^ did not cross the membrane in the absence of Tween-20 (**Figure S7**), (ii) the Cal-520 fluorescence is insensitive to Tween-20 (**Figure S8**), and (iii) the dye operates in Tween-20 rich solutions (**Figure S8**), providing confidence that the observed fluorescence enhancements are due to Tween-20 induced membrane permeabilization and solution exchange. Similar intensity enhancements were observed at 4°C and 37°C reflecting a Ca^2+^ influx at the end of the titration of 98 % and 96 %, respectively (**Figure 2D**), though it is of particular interest to note that Hill models applied to the Cal-520 intensity enhancement data revealed half-maximal concentration constants substantially lower than those observed under the ensemble FRET measurements (**Table S2**), pointing towards a situation where the concentration requirement to achieve membrane permeabilization is lower than that required to fully separate the lipophilic probes in the FRET assay.

To establish whether the observed fluorescence signals were correlated with changes in vesicle morphology, we next performed DLS as a means to assess the hydrodynamic diameters (d_H_) of vesicles in the ensemble. As reported by us and others(23, 24, 40, 41), DLS is highly sensitive to the diffusion of vesicles in solution, making it particularly attractive to quantify the particle size distribution as a function of Tween-20. In the absence of Tween-20, the diameter of freshly prepared vesicles was found to be 194 ± 2 nm with a polydispersity index of ∼0.47 (**Figure 2E, Figure S9**). In the presence of low concentrations (0 – 15 μM) of Tween-20, the correlation curves progressively shifted towards longer lag times, consistent with an increase in diffusion coefficient and ∼75 % increase in vesicle size. This observation was attributed to vesicle expansion, fusion or the combination of both given the mean polydispersity index increased towards 20 μM Tween-20 before then reducing in magnitude likely as a consequence of micellization (**Figure S9**).

To minimize the possibility of vesicle fusion and explicitly test whether expansion was taking place, we next moved to investigate the hydrodynamic diameters using FCS. Unlike DLS, FCS is collectively used on systems in which the absolute concentration of the fluorescent species is in the sub-nanomolar range, hence the particles interact negligibly with each other. The technique is thus an ideal tool for testing for structural variations in single freely-diffusing vesicles. As illustrated in **Figure 2F and Figure S10**, the diffusion coefficients and d_H_ of POPC vesicles prepared with 0.1 % DiI were recovered from the correlation functions obtained from vesicles freely-diffusing through a confocal volume. In the absence of detergent, we recorded a hydrodynamic diameter of 208 ± 12 nm and diffusion coefficient of 2.1 ± 0.1 μm^2^s^-1^. Assuming spherical vesicles, the mean diameter was found to increase by ∼20 % in the presence of 0.06 mM Tween-20, whereas the mean diffusion coefficient reduced to ∼84 % of its initial value, suggestive of substantial structural remodelling assigned to expansion within single in-tact vesicles. It is noteworthy that the relative increase in hydrodynamic diameter and translational diffusion time is comparable to those of similar size and composition in the presence of low concentrations of Triton-X 100, which we interpreted as evidence that Tween-20 leads to a similar degree of vesicle perturbation at the detergent cmc.

### Tween-20 Induces Structural Remodelling of Surface-Tethered POPC Vesicles

To further explore the structural changes, we next performed single-vesicle FRET imaging by using a custom-built wide-field, objective-based total internal reflection fluorescence microscope that enables the simultaneous and parallel imaging of DiI and DiD emission(24, 42). Using this approach, the mean FRET efficiencies of single surface-immobilized vesicles labelled with 0.1% DiI and 0.1 % DiD were monitored upon addition of Tween-20 at 21°C. Surface-immobilization was achieved by incorporating 1% biotinylated lipids into the vesicle structure to facilitate their immobilization to a glass coverslip via biotin-NeutrAvidin interactions (**Figure 3A**). An oxygen scavenger cocktail consisting of glucose oxidase, catalase and Trolox was also added to the imaging buffer. Trolox, a vitamin E analogue, minimizes photoblinking, while the glucose oxidase and catalase system reduces molecular oxygen and photobleaching by oxidizing glucose(43, 44). In the absence of Tween-20, ∼150-200 vesicles per 25 × 50 μm field of view were imaged, representing surface-tethered vesicles separated on average by a nearest-neighbour separation distance of ∼1 μm (**Figure 3B**). When imaged under low excitation power (8.2 mWcm^-2^), the donor, acceptor and FRET trajectories remained stable over the duration of the measurement, pointing toward high photostability over a 50 s time-window (**Figure 3C**). Because of efficient FRET between DiI and DiD within in-tact vesicles, strong fluorescence emission is observed across both donor and acceptor detection channels in the absence of Tween-20. Changes to the donor and acceptor fluorescence intensities after addition of Tween-20 were then recorded for several hundred single, spatially separated vesicles. In this case we observed a substantial decrease in the acceptor emission and a corresponding increase in the relative donor emission because of reduced FRET between the dyes (**Figure 3C**). It is noteworthy that upon Tween-20 addition at 0.06 mM and 0.12 mM, the mean number of fluorescent spots per field of view, the mean total emission intensity defined as <I_T_> = <I_D_+I_A_> (where I_D_ and I_A_ are the donor and acceptor emission intensities respectively), and mean nearest-neighbour vesicle separation distance, <d> remained largely invariant (**Figure 3D**), indicating the presence of in-tact vesicle structures. At higher detergent concentrations, a ∼2-fold increase in the nearest neighbour separation distance and corresponding decrease in <I_T_> was observed, indicating removal of lipid material from the surface. All fluorescence/FRET trajectories were acquired over a 50 s time window after Tween-20 addition using 50 ms exposure. The FRET efficiency drop observed across individual vesicles at low Tween-20 concentrations (**Figure 3E**) could not therefore be attributed to lipid loss or partial vesicle detachment because the corresponding total fluorescence intensity and nearest-neighbour separation remained unchanged. Instead, this observation points towards a substantial structural change, namely swelling, taking place within individual surface-immobilized vesicles, with minimal loss of lipid material to the solution. Notably, this result is opposed to previous observations where Tween-20 had no effect on the modal size distribution of human-derived extracellular vesicles (EVs) (45). An explanation for this discrepancy rests in the lipid composition; while in this work we used highly-controllable model-membranes composed primarily of POPC lipids, most studies indicate that EVs are enriched in cholesterol compared with the total lipid pool(46). As we previously demonstrated, PC vesicles composed of modest cholesterol content can lead to a substantial resistance to detergent induced solubilization, likely due to cholesterol obstructing the initial step of detergent molecules inserting into the lipid bilayer(23). In strong contrast to EVs, the vast majority of vesicles studies here (> 95 %) showed the swelling behaviour. When 0.18 mM, 0.24 mM and 0.3 mM Tween-20 solutions were added to the immobilized vesicle layer, the nearest neighbour separation distance progressively increased to 1.2 ± 0.1 μm, 2.7 ± 0.1 μm and 2.6 ± 0.2 μm, respectively, corresponding to a reduction in the number of vesicles per field-of-view which we assigned to loss of material and the removal of a fraction of the vesicle population (**Figure 3D**). However, with increasing levels of Tween-20 those that remained on the surface displayed, a stepwise shift towards lower FRET efficiency signatures indicative of further expansion and remodelling.

**Fig. 3.**
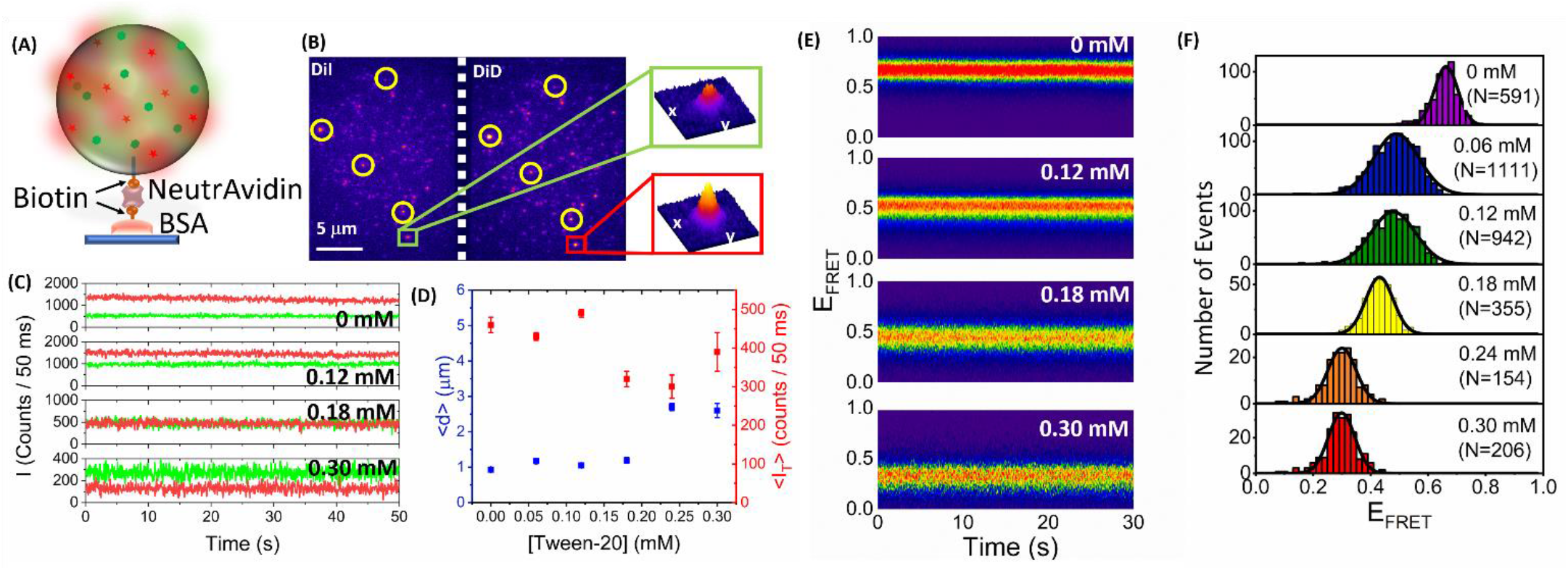
Single vesicle FRET imaging of surface-tethered POPC vesicles. (A) Schematic representation of the methodology used for surface-tethering DiI and DiD loaded vesicles. Large unilamellar vesicles composed of 98.8% POPC, 0.1 % DiI, 0.1 % DiD and 1 % Biotin-PE were attached to a glass coverslip via BSA using biotin-NeutrAvidin chemistry. (B) Representative wide-field TIRF image obtained from DiI and DiD loaded POPC vesicles (λ_ex_ = 532 nm) in the absence of Tween-20, showing representative acceptor signals colocalized with their corresponding donors (yellow circles). Exposure time = 50 ms. Insets: 3D surface intensity plots of DiI and DiD emission from a single surface-immobilized vesicle. The intensity axes are scaled to a dynamic range of 0-10 counts to provide visual contrast and facilitate comparison. (C) Representative fluorescence emission intensity profiles of individual POPC vesicles labelled with DiI (green) and DiD (red) in the absence and presence of 0.12 mM, 0.18 mM and 0.3 mM Tween-20. (D) Corresponding variation in mean nearest-neighbour separation distance, <d>, and mean total spot intensity, <I_T_ > obtained as a function of [Tween-20]. (E) Contour plots of the time-evolution of the FRET population as a function of Tween-20. Each contour plot was produced by superimposing individual FRET trajectories and contours are plotted from blue (lowest population) to red (highest population). (F) Histograms of the mean apparent FRET efficiency, E_FRET_, obtained from single immobilized vesicles after incubation of Tween-20 across a broad concentration range. Solid black lines represent Gaussian fits to the experimental data. Solution conditions: 50 mM Tris, 6 % (W/V) D-(+)-glucose, 1 mM Trolox, 6.25 μM glucose oxidase, 0.2 μM catalase, pH 8.

From the single-vesicle images and fluorescence trajectories, histograms of the mean energy transfer efficiencies were generated from several hundred single vesicles per condition, allowing for discrete conformation changes to be identified (**Figure 3F**). Under all conditions tested, the FRET efficiency distributions displayed Gaussian behaviour, and in the absence of Tween-20, a peak FRET efficiency of 0.66 ± 0.08 was observed, corresponding to in-tact vesicles where the dyes are spatially separated close to their Förster radius. With increasing Tween-20 concentrations towards 0.18 mM, the FRET population decreased in a stepwise manner towards 0.43 ± 0.05, indicative of a ∼35 % increase in the mean dye-pair separation distance. At higher concentrations (0.24 mM and 0.30 mM), the peak position shifted further to 0.30 ± 0.09. Overall, the reduction in the mean apparent FRET signature agrees well with the conformational changes reported by QCM-D, DLS and FCS and confirms Tween-20 induces the structural remodelling of single surface-tethered and intact vesicles.

To further probe the structure and size distribution of single vesicles in response to Tween-20, we also employed low voltage scanning electron microscopy (SEM) as previously described(47). Images acquired by SEM revealed that in the absence of Tween-20, the vesicles were predominantly spherical (circularity = 0.68 ± 0.01) in morphology with a mean diameter of 72 ± 3 nm (**Figure 4a**) (N = 137). In the presence of Tween-20 at concentrations similar to those used in the single-vesicle assay, the mean circularity was similar (0.67 ± 0.01) (**Figure 4b, c**), however, the size distribution substantially broadened (**Figure 4d**). While our SEM sample preparation utilized only a thin (5 nm) conductive layer, which as previous studies indicate does not substantially alter the imaging result(48), we note that the intrinsic requirement to dehydrate the vesicles may be an explanation for why the observed particle size in the absence of detergent is lower than those reported by FCS and DLS. Nevertheless, our observations of spherical morphologies are in line with similar studies by others(49, 50) and taken in conjunction with our fluorescence and QCM-D data, the results are also broadly supportive of the Tween-20 induced swelling of single intact vesicles.

**Fig. 4.**
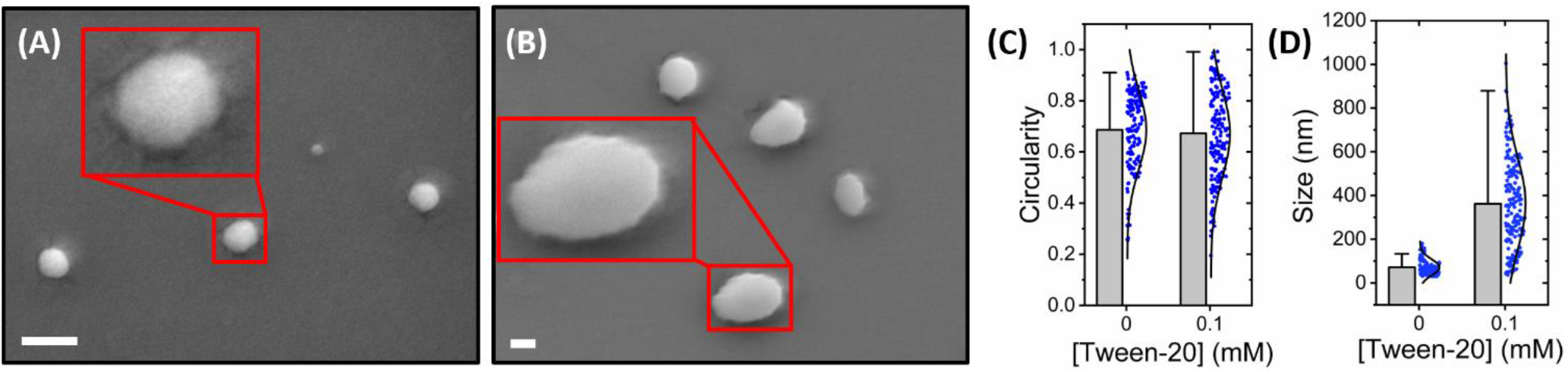
SEM analysis of single lipid vesicles. Representative electron micrographs of vesicles (A) in the absence and (B) in the presence of 0.1 mM Tween-20. Scale bars = 100 nm. Also shown are comparative bar plots summarizing the relative variation in (C) circularity and (D) particle size for vesicles in the absence (N = 137) and presence (N = 176) of Tween-20.

### Calcium Influx is Induced upon Interaction of Tween-20 with surface-tethered POPC Vesicles

The QCM-D, DLS, FCS and single-vesicle FRET measurements all point towards the global remodelling of the vesicle structure in response to Tween-20. However, whether the interaction also involved content leakage via membrane permeabilization remained an open question. To assess this, we moved to investigate membrane integrity during interaction with Tween-20 by monitoring the influx of calcium into single surface-tethered POPC vesicles loaded with Cal-520. We incubated the immobilized vesicles in 50 mM Tris, pH 8.0 containing Ca^2+^ at a concentration of 10 mM and imaged them under TIRF conditions in the absence and presence of Tween-20. Addition of Tween-20 to a sample containing surface-tethered, Cal-520 loaded POPC vesicles then resulted in an increase in the Cal-520 fluorescence signal as a consequence of membrane permeabilization, as schematically illustrated in **Figure 5A**. Typically, we imaged 10 fields of view for each condition, allowing us to quantify the fluorescence emission intensities of several thousand vesicles, before adding buffer solutions rich in Tween-20 and Ca^2+^. If Ca^2+^ gained entry into a vesicle, an increase in fluorescence intensity was detected, leading to substantial variation in the overall intensity distribution. In the absence of Tween-20 and Ca^2+^, ∼100-150 vesicles per field of view were identified (**Figure 5B**) and the intensity distribution displayed lognormal behaviour with a peak of ∼85 counts/100 ms, which is to be expected for diffusing molecules(51) (**Figure 5E**). When 10 mM Ca^2+^ was flushed over the vesicles, the number of spots remained largely unchanged (**Figure 5C**), and the intensity distribution was broadly similar (**Figure 5E**) peaking at ∼91 counts/100 ms, indicating minimal levels of Ca^2+^ influx into the in-tact vesicles. However, in the presence of 0.01-0.04 mM Tween-20, we observed a clearly discernible difference, with the vesicle spots appearing both progressively larger and brighter (**Figure 5D**). This is more clearly observed in the lognormal distribution of spot intensities, which show a transition from low-to-high (> 200 counts/100 ms) values even at modest concentrations of Tween-20 (**Figure 5E, Figure S11**). At higher concentrations of 0.05 mM and 0.06 mM Tween-20, we observed a shift in the peak position towards lower intensity values, which we hypothesise could be due to Cal-520 leakage. It is worth noting that in all cases of [Tween-20], a 2-fold or greater increase in the single vesicle intensity was observed, in line with our ensemble measurements and those observed by others(33). Importantly, the fluorescence signals before and after addition of Tween-20 were stable (**Figure S11**) and the number of surface-immobilized vesicles remained invariant as Tween-20 was flushed across the surface. Taken together, the combined data implies that the interaction between Tween-20 and highly-curved POPC vesicles not only leads to substantial vesicle remodelling, but that the solubilization mechanism also involves membrane permeabilization and ultimately solution exchange between the vesicles and the local environment. Despite only moderate changes in the Cal-520 emission intensity, this analysis suggests particularly substantial Tween-20 induced permeabilization, given a doubling of signal intensity was also observed in the presence of ionomycin.

**Fig. 5.**
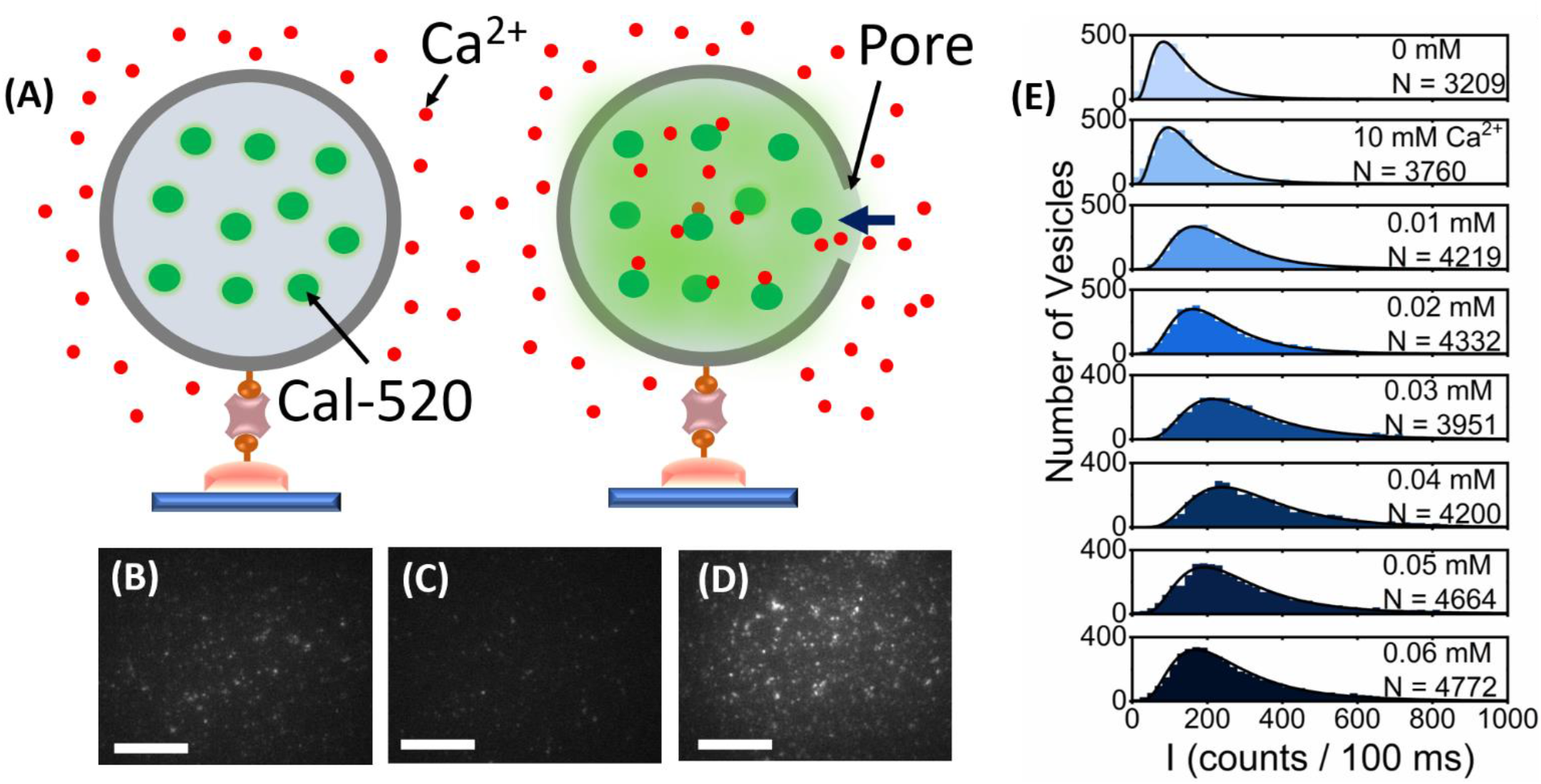
Changes in encapsulated Cal-520 fluorescence intensity upon addition of Tween-20. (A) Schematic illustration of the assay. 200 nm sized POPC vesicles encapsulating Cal-520 are placed in buffer solution containing Ca^2+^ (left panel). Membrane permeabilization after addition of Tween-20 results in Ca^2+^ influx and fluorescence intensity enhancement of Cal-520. Representative wide-field TIRF images of Cal-520-loaded vesicles (λ_ex_ = 488 nm) in (B) imaging buffer (50 mM Tris, pH 8), (C) imaging buffer including 10 mM Ca^2+^ and (D) imaging buffer including 10 mM Ca^2+^ and 0.01 mM Tween-20. (E) Intensity population histograms obtained from N > 3000 Cal-520-loaded vesicles in imaging buffer (top panel), imaging buffer including 10 mM Ca^2+^, and imaging buffer including 10 mM Ca^2+^ and Tween-20 (lower panels). The solid black lines represent lognormal fits to each of the identified peaks.

## Discussion

One of the most remarkable outcomes of this study is the multi-step nature of the Tween-20 interaction with POPC vesicles prior to lysis. This broadly involves (1) the deposition of detergent molecules onto the membrane surface, (2) conformational restructuring that can be assigned to vesicle expansion and (3) membrane permeabilization that leads to solution exchange. Notably, these events cannot be identified and distinguished using a single stand-alone technique, but rather they have emerged by utilizing a highly interdisciplinary toolkit comprising QCM-D, ensemble and time-resolved fluorescence measurements, DLS, FCS, and single-vesicle imaging. Owing to its bulky headgroup and pliable hydrocarbon chain, Tween-20 can be modelled as a cone with positive spontaneous curvature, and thus we attribute the observed structural changes to the incorporation of surfactant molecules into the lipid bilayer. Moreover, large unilamellar vesicles exhibit high radii of curvature which we hypothesise destabilizes during the interaction. This then likely leads to a reduction in membrane line tension, which facilitates the global deformability of the bilayer, and promotes both expansion and the formation of pores through which Ca^2+^ can enter (20, 52). Indeed, this assertion is supported by additional fluorescence lifetime measurements involving the interaction with POPC vesicles coated in the membrane tension probe FliptR (**Figure S12**). As previously reported(53), the probe lifetime depends linearly on the membrane tension, and in our case, a reduction in lifetime from 3.35 ± 0.02 ns in the absence of Tween-20, to 2.77 ± 0.03 ns at ∼10x the cmc was observed, indicative of probe de-planarization. Even if only a single calcium ion then enters the vesicle, the local Ca^2+^ concentration increases by several hundred nM, which is readily detectable via the Cal-520 signal enhancement assay(33). It is of interest to note that this scenario, while consistent with recent work suggesting Tween-20 induces bulging of live cells(4), differs from those observed using extracellular vesicles, where the vesicle concentration reduced gradually with increasing Tween-20, but without variation in particle size(45, 54). In the latter case, the concentration of highly curved extracellular vesicles (EVs) with a similar size distribution to those used here, was strongly modulated by Tween-20, especially at detergent concentrations above the cmc, with the observed concentration reduction attributed to either large vesicle aggregates breaking apart or the lysis of single EVs. This difference can be qualitatively explained by taking the membrane composition into account, and we speculate that the presence of high cholesterol content in EVs likely obstructs the initial detergent-membrane interaction. In contrast, membranes with low or no cholesterol content, such as the vesicles used in this work, facilitate the insertion of Tween-20, triggering the observed expansion, permeabilization, and solution exchange. Indeed a fundamental requirement of membrane leakage and structural remodelling is membrane insertion, suggesting that the detergent may penetrate deeply into the POPC membranes to trigger the observed effects. The observation of a multi-step solubilization process is further supported by previous optical microscopy experiments involving ∼10 μm sized vesicles composed of 1,2-dioleoyl-*sn-*glycero-3-phosphocholine (DOPC) where Tween-20 induced long-lived pores across a measurement timescale of ∼125 s(52). In this study, a concentration gradient resulted in transient, but long-lived pores capable of inducing solution exchange and was explained by a substantial reduction in the membrane line tension. A direct quantitative comparison between our findings on ∼200 nm sized vesicles and the giant vesicles used in this work is not straightforward due to variations in vesicle size and composition, the use of a concentration gradient and the fact that optical imaging only provides access to a cross-section of the focal plane. In contrast, our approach involves the use of immobilized vesicles in a microfluidic flowcell where a steady-state detergent concentration can be rapidly reached, and our single-vesicle approach utilizes the mean FRET signature from across the three-dimensional volume of the vesicle. Nevertheless, it is clear that giant vesicles and the smaller sub-micron vesicles studied here have common solubilization attributes including similar concentration requirements and the presence of a solution exchange step attributed to the formation of pores. Thus, our work on highly curved vesicles are complementary of previous studies on giant vesicles and points to common structural remodelling prior to complete solubilization.

An additional aspect of our results that deserves attention is that a low concentration of Tween-20, below the cmc, produced substantial structural alterations. One possible explanation for this is the formation of discrete membrane regions with a high local detergent density that acts as a nucleation site for the observed rearrangements. This is partially supported by the application of a mass-action model to the FRET efficiency curves, which reveal limited compatibility of the surfactant in the membrane in terms of a Flory-Huggins parameter. Similar interpretations have already been adopted in the context of protein-induced disruption of ∼100 nm sized 1,2-dioleoyl-sn-glycero-3-phospho-L-serine (DOPS) and 1,2-dioleoyl-sn-glycero-3-phospho-(1′-rac-glycerol)) (DOPG) vesicles(55) and is broadly supported by our FCS and single-vesicle FRET assays where discrete changes to the observed hydrodynamic radii and FRET efficiency respectively, representing discrete conformational changes, was observed to occur in a stepwise manner as the Tween-20 concentration progressively increased towards the cmc. Indeed our wide-field single-vesicle FRET and Cal-520 imaging approaches enabled both expansion and solution exchange within in-tact vesicles to be monitored at lipid: detergent ratios of ∼ 2×10^3^, suggesting that the vesicle structural rearrangements and vesicle leakage are triggered by < 100 Tween-20 molecules per 200 nm sized vesicle. From inspection of the fluorescence traces, the observed expansion and solution exchange occurs without complete removal of lipid material from the surface.

In summary, we have established that the combination of solution and surface-based ensemble measurements, combined with ultrasensitive single-vesicle spectroscopy approaches, reveal precise molecular level events that underpin how Tween-20 disrupts lipid vesicles in vitro. By using a combination of QCM-D, ensemble and picosecond time-resolved fluorescence approaches, DLS, FCS and wide-field single-vesicle imaging, we have established that Tween-20 dynamically alters the structure and integrity of both freely-diffusing and surface-immobilized model-membrane vesicles. We show that Tween-20 induces the solubilization of vesicles composed of POPC lipids via a mechanism involving an initial mass gain that precedes a structural remodelling event and permeabilization that leads to content exchange. Taken together, these observations provide new mechanistic insights for how solubilizing detergents perturb and damage highly curved lipid membranes, and may be directly relevant to a number of biotechnological applications where conformational control of the membrane is vital. We also expect that the developed approaches will find vast utility in revealing precise, nanometre sized structural changes taking place in lipid vesicles in response to a wide variety of perturbative agents, including membrane disrupting proteins, surfactants and anti-viral agents.

## Methods

### Materials

Lipids suspended in chloroform (1-palmitoyl-2-oleoyl-glycero-3-phosphocholine (POPC), 1-oleoyl-2-(12-biotinyl(aminododecanoyl))-sn-glycero-3-phosphoethanolamine (biotin-PE) and Tween-20 were purchased from Sigma Aldrich. Tween-20 was suspended in 50 mM Tris buffer (pH 8) prior to each use. Lipophilic membrane stains (1′-Dioctadecyl-3,3,3′,3′-Tetramethylindocarbocyanine Perchlorate (DiI) and 1,1′-Dioctadecyl-3,3,3′,3′-Tetramethylindodicarbocyanine, 4-Chlorobenzenesulfonate Salt) (DiD)) were purchased from ThermoFisher Scientific. Cal-520, sodium salt was purchased from Stratech and freshly suspended in MilliQ water prior to use. POPC, Biotin-PE and Cal-520 solutions were stored at −20°C, whereas DiI and DiD solutions were stored at 4°C prior to use. All samples were used without additional purification.

### Preparation of vesicles incorporating the DiI and DiD FRET sensor

Large unilamellar vesicles composed of 0.1 mol % DiI, 0.1 mol % DiD, 1 % biotin-PE and 98.8 mol % POPC were prepared as previously described(23, 24). Briefly, lipids, DiI and Did were mixed in chloroform before the solvent was evaporated. The dried lipid film was then resuspended in 50 mM Tris buffer (pH 8) and vortexed. Large unilamellar vesicles were then prepared by the extrusion method, whereby the solutions were extruded at least 21 times through a polycarbonate membrane filter with size cut off of 200 nm. The polydispersity index, as measured by DLS, was 0.478 ± 0.04. On the basis of monodisperse LUVs of 200nm diameter, and reports confirming that both DiI and DiD have excellent membrane-incorporation efficiency and retention, while also minimally perturbing membrane morphology(56), the average number of fluorophores incorporated per vesicle,<N>, was estimated via 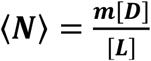, where m is the average number of lipids per vesicle, [D] is the total dye concentration and [L] is the total lipid concentration(57). The lipid head group was estimated to have a surface area of 0.92 nm^2^, resulting in a value of m ∼ 2 × 10^5^ lipids per vesicle, and <N_DiI_> = <N_DiD_> ∼ 200. Using a similar approach, POPC vesicles coated in FliptR tension probe were prepared via extrusion using a dye concentration of 4 μM.

### Preparation and purification of Cal-520 loaded vesicles

Cal-520 filled vesicles were prepared as previously described(33). Briefly, POPC and Biotin-PE chloroform stock solutions were mixed such that the POPC: Biotin-PE molar ratio was 100: 1. The chloroform was then removed and the samples were dissolved and vortexed in 50 mM Tris buffer (pH 8) containing 100 μM Cal-520. The solutions were then extruded at least 21 times through a polycarbonate membrane filter with pore size of 200 nm. Vesicle solutions were filtered using a PD-10 desalting column (Sigma Aldrich) and size-exclusion chromatography was performed using an AKTA pure system (GE Healthcare) in order to separate free Cal-520 dye molecules from loaded vesicles.

### Quartz Crystal Microbalance with Dissipation (QCM-D) Monitoring

Time-dependent changes in frequency and dissipation were recorded using a Q-Sense E4 (Biolin Scientific) system. SiO_2_-coated sensors (Biolin Scientific) with a fundamental frequency of 5 MHz were treated with UV/ Ozone for 10 minutes, then sonicated in 2 % Hellamanex III for 10 minutes and ultrapure Milli-Q water for 20 minutes and dried under N_2_ flow. The sensors were then treated with UV/ Ozone for a further 30 minutes, immersed in ethanol and dried with N_2_ flow prior to use. Each sensor was pre-functionalized with an amine monolayer by immersion in 4% v/v ATES/IPA solution for 16 hours, followed by washing with IPA and dried with N_2_ before installation in the flow modules. The sensors were then flushed with 50 mM Tris buffer (pH 8) at 20 μL / min until a stable baseline was reached. At the start of each experiment, the sensor surfaces were coated with 0.1 mg/mL biotinylated BSA (Sigma Aldrich) and 1 mg/mL BSA (Sigma Aldrich) dissolved in 50 mM Tris buffer (pH 8). When saturation was reached, the sensors were rinsed with buffer to remove unbound molecules, before NeutrAvidin (ThermoFisher Scientific) was added at 0.2 mg/mL in 50 mM Tris buffer (pH 8) until saturation was reached and a further rinse step with buffer was performed. Vesicles containing 1 mol % biotin-PE and 99% POPC were then added to the surface using a final lipid concentration of 25 μg/mL in 50 mM Tris buffer until saturation was reached. This was followed by a further rinse step and the insertion of Tween-20 detergent solutions in 50 mM Tris buffer (pH 8) at the specified concentrations.

### Steady State Fluorescence and FRET Spectroscopy

Fluorescence emission spectra were measured using continuous wave excitation under magic angle conditions with a HORIBA Fluoromax-4 spectrophotometer. Spectra from vesicles containing DiI and DiD were recorded using an excitation wavelength of 520 nm, whereas an excitation wavelength of 493 nm was used for measuring Cal-520 filled vesicles. Apparent FRET efficiencies, E, were determined via E = I_A_/(I_A_+I_D_) where I_D_ and I_A_ represent the fluorescence emission intensities of DiI and DiD, respectively. FRET efficiency plots were fitted using *E* = *A* + *Bϕ*, where *A* and *B* are the measured FRET efficiencies at 0 mM Tween-20 and at the end of the titration, respectively, and 0 ⩽ *ϕ* ⩽ 1 is the volume fraction of lipids in the membrane. As molecular models, we used the Hill model, *ϕ* = 1/(1 + ([*Tween* − 20]/*k*)^*n*^), with k the half-maximal concentration and *n* the Hill coefficient, and we used a mass-action model for mixed micellisation *kexp*(−*ϕ*^2^*χ*)([*Tween* − 20]/(1 − *ϕ*) − [*lipid*]/*ϕ*) = 1, where K is an equilibrium constant and *χ* is the Flory-Huggins interaction parameter (**Supplementary Text 1**). The relative influx, *φ*, of calcium into vesicles upon o addition of Tween-20 was determined via 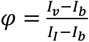 where I_v_ is the Cal-520 intensity after addition of Tween-20, I_b_ is the Cal-520 intensity in the absence of Tween-20 and I_I_ is Cal-520 intensity after addition of 1mg/mL ionomycin(33, 58). The data points shown in **Figure 2** represent the mean and standard error of the mean from 3 individual experimental runs and all experiments were performed in 50 mM Tris buffer (pH 8) with a final POPC concentration of 25 μM.

### Fluorescence Lifetime Measurements

Time-resolved fluorescence spectroscopy on DiI and DiD loaded vesicles was performed using a FluoTime300 time-correlated single photon counting spectrophotometer equipped with a hybrid PMT detector (PMYA Hybrid 07, Picoquant). Time-resolved fluorescence decays were measured under magic angle conditions using pulsed excitation at 532 nm with a repetition rate of 50 MHz (LDH-D-FA-530L, Picoquant). Fluorescence emission decays at 565 nm, corresponding to the peak of DiI emission, were collected until 10^4^ counts accumulated at the decay maximum. Fluorescence decay curves were fitted by iterative re-convolution of the instrument response function and the observed fluorescence decay using a multi-exponential decay function of the form 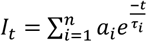 where I_t_ is the intensity at time, t, normalized to the intensity at t = 0, and t_i_ and a_i_ represent the fluorescence lifetime and fractional amplitude of the i’th decay component. The quality of the fit was judged based on the convergence of the reduced chi-squared. All experiments were performed in 50 mM Tris buffer (pH 8) with a final POPC concentration of 25 μM. In the context of FliptR measurements, fluorescence lifetime decays were acquired at 600 nm using a pulsed 485 nm laser (LDH-P-C-485, Picoquant) with a repetition rate of 20 MHz.

### Dynamic Light Scattering

Vesicle size distributions in the absence and presence of Tween-20 were measured using a Zetasizer μV DLS system (Malvern Instruments). A final POPC concentration of 50 μM in 50 mM Tris buffer (pH 8) was used in all experiments. Briefly, the Brownian motion of vesicles in suspension scattered laser light at 632.8 nm, and correlation of the intensity fluctuations yields the diffusion coefficient and hydrodynamic radius via the Stokes-Einstein relationship as previously described(17, 27). All DLS data were collected using 178° backward scattering and averaged over three experimental runs. The refractive index of the dispersion medium was 1.33. All sizes discussed are in terms of spherical hydrodynamic radii, and all DLS analyses are reported in terms of intensity distribution.

### Fluorescence Correlation Spectroscopy

Fluorescence correlation spectroscopy (FCS) measurements were performed on a Zeiss LSM 880 microscope, using a GaAsP detector as previously described(59). Samples composed of 99.9 mol % POPC and 0.1 mol % DiI were pipetted onto microscope slides, sealed with silica and a 1.5 coverslip. Samples were excited with a 514 nm excitation line with typical powers of 4 μW as measured at the sample plane. Final concentrations of POPC and DiI in 50 mM Tris buffer (pH 8) were 70 μM and 80 nM, respectively. The confocal volume was measured using a calibration sample of 6 nM Rhodamine-6G at 21°C and constraining the diffusion coefficient to be ∼400 μm^2^s^-1^. Autocorrelation curves, G(τ) were fitted to an expression of the form 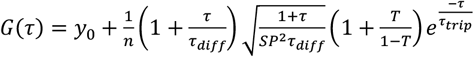 where y_0_, n, τ, τ_diff_, SP, T and τ_trip_ are offset, effective number of particles in confocal volume, lag time, residence time in confocal volume, structure parameter, fraction of particles in triplet state and residence time in triplet state respectively. Diffusion coefficients, D, were then determined by 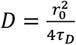, where r is the spot width (∼0.238 μm).

### Single Vesicle FRET Spectroscopy

Microfluidic flow cells were constructed as described previously(42) and coated with 0.1 mg/mL BSA-Biotin, 1 mg/mL BSA and 0.2 mg/mL Neutravdin as described for our QCM-D monitoring approach. Biotinylated POPC vesicles containing 0.1 mol % DiI and 0.1 mol% DiD were then added to a final concentration of 70 μg/mL in imaging buffer (50 mM Tris, 6 % (W/V) D-(+)-glucose containing 1 mM Trolox and 6.25 μM glucose oxidase and 0.2 μM catalase) and incubated for 15 minutes at room temperature to achieve a surface coverage of ∼150-200 vesicles per 50 × 50 μm field of view. Unbound vesicles were then removed by washing the flowcell with imaging buffer. Bespoke TIRF microscopy was then performed on an inverted microscope (Nikon Eclipse Ti) containing a CFI Apo TIRF 100 x NA 1.49 oil-immersion objective lens (Nikon) and illumination from a TEM_00_ 532 nm line (Obis, Coherent). Emission was separated from the excitation line via a dichroic and emission filter mounted beneath the lens turret (Chroma 59907-ET-532/640). DiI and DiD emission was then spatially separated using a DualView image splitter containing a dichroic filter (T640LPXR, Chroma) and band pass filters (ET585/65M and ET700/75M, Chroma) and imaged in parallel on a back-illuminated Prime 95b CMOS camera cooled to −30°C (Photometrics). After each addition of Tween-20 in imaging buffer, movies were acquired with 50 ms exposure time. Recorded images were then analysed in MATLAB (R2019a) using iSMS single-molecule FRET microscopy software(60). Briefly, the donor and acceptor emission channels were aligned, and background-corrected DiI and DiD emission trajectories were obtained by integration of the intensity within the area containing the vesicle signal for each time point. Apparent FRET efficiencies were calculated as described previously, and related to the mean distance between probes, R, via 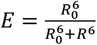, where R_o_ = 5.3 nm is the Förster radius.

### Scanning Electron Microscopy

Scanning electron microscopy was performed using a JEOL JSM 7800-F system operating at 5kV. Vesicles were prepared in 50 mM Tris (pH 8) containing Tween-20 at the specified concentrations, diluted ∼10 x in deionized water and vortexed. A 30 mL volume of the vesicle solution was then added to a silicon substrate and the solution evaporated. The substrate was then covered with a 5 nm Pt/Pd layer to avoid any charging effects or disruption of the vesicles in the microscope. Vesicle diameters were then determined using ImageJ, where automated analysis of black-and-white binary images enabled separation of regions of white pixels (the vesicles) against a dark background. Vesicle circularity was measured via 4π(A/p^2^), where A is the observed area and p is the perimeter. In this case, a circularity value of 1 indicates a perfect circle, whereas a circularity value approaching 0 indicates an increasingly elongated polygon.

### Single Vesicle Measurements of Ca^2+^ Influx

Biotinylated POPC vesicles encapsulating Cal-520 at a concentration of 200 μM were prepared and immobilized as previously described. Tethered vesicles were then incubated with imaging buffer containing 10 mM Ca^2+^ and imaged via TIRF using 488 nm excitation. Cal-520 fluorescence was separated from incident excitation using a dichroic mirror of edge wavelength 495 nm and emission bandpass filter (502-548 nm) and imaged using objective-based TIRF. Next, Ca^2+^ rich imaging buffer containing Tween-20 at concentrations specified in the main text were added to the flowcells and images of Ca^2+^ saturated vesicles were acquired with an exposure time of 100 ms.

## Data availability

All data, procedures and protocols are provided in the manuscript and Supporting Information.

**The authors declare no competing interests**.

## ACKNOWLEDGMENTS

This work was supported by Alzheimer’s Research UK (RF2019-A-001) and EPSRC (EP/V034030/1, EP/N031431/1, EP/T002166/1 and EP/N031431/1). We thank Prof. Daniella Barillá (Department of Biology, University of York) for use of DLS instrumentation, Dr Jared Cartwright (Department of Biology, University of York) for use of size exclusion chromatography apparatus and Prof. Thomas Krauss (University of York) for use of SEM facilities. We also thank the Bioscience Technology Facility for use of FCS and fluorescence spectroscopy facilities (University of York, UK) and Prof. Marco Fritzche (University of Oxford, UK) for the generous donation of FliptR.

## Author contributions

S. D. Q. designed research. L. D., S. G., D. C. and L. M performed research; S. J. and M. L. contributed reagents. S. J., C. S., and M. C. L. contributed analytic tools; L. D., S. G., L. M., C. S., S. J., M. C. L. and S. D. Q. analyzed data; and S. D. Q. wrote the paper with the help of the other co-authors.

## References

1. Hjerten S, Johansson KE. Selective solubilization with Tween-20 of membrane proteins from Acholeplasma laidlawaii. Biochim Biophys Acta. 1972;288(2):312–25.

2. Schuck S, Honsho M, Ekroos K, Shevchenko A, Simons K. Resistance of cell membranes to different detergents. Proc Natl Acad Sci U S A. 2003;100(10):5795–800.

3. Mayo DR, Beckwith WH, 3rd. Inactivation of West Nile virus during serologic testing and transport. J Clin Microbiol. 2002;40(8):3044–6.

4. Hua T, Zhang X, Tang B, Chang C, Liu G, Feng L, et al. Tween-20 transiently changes the surface morphology of PK-15 cells and improves PCV2 infection. BMC Vet Res. 2018;14(1):138.

5. Cipolla D, Wu H, Gonda I, Eastman S, Redelmeier T, Chan HK. Modifying the release properties of liposomes toward personalized medicine. J Pharm Sci. 2014;103(6):1851–62.

6. Sahoo RK, Biswas N, Guha A, Sahoo N, Kuotsu K. Nonionic surfactant vesicles in ocular delivery: innovative approaches and perspectives. Biomed Res Int. 2014;2014:263604.

7. Jiang R, Wu X, Xiao Y, Kong D, Li Y, Wang H. Tween-20 regulate the function and structure of transmembrane proteins of Bacillus cereus: Promoting transmembrane transport of fluoranthene. J Hazard Mater. 2021;403:123707.

8. Chan YHM, Boxer SG. Model membrane systems and their applications. Curr Opin Chem Biol. 2007;11(6):581–7.

9. Nomura F, Nagata M, Inaba T, Hiramatsu H, Hotani H, Takiguchi K. Capabilities of liposomes for topological transformation. Proc Natl Acad Sci U S A. 2001;98(5):2340–5.

10. Hamada T, Hagihara H, Morita M, Vestergaard MC, Tsujino Y, Takagi M. Physicochemical Profiling of Surfactant-Induced Membrane Dynamics in a Cell-Sized Liposome. J Phys Chem Lett. 2012;3(3):430–5.

11. Chabanon M, Rangamani P. Solubilization kinetics determines the pulsatory dynamics of lipid vesicles exposed to surfactant. Biochim Biophys Acta Biomembr. 2018;1860(10):2032–41.

12. Karatekin E, Sandre O, Brochard-Wyart F. Transient pores in vesicles. Polym Int. 2003;52(4):486–93.

13. Hamada T, Hirabayashi Y, Ohta T, Takagi M. Rhythmic pore dynamics in a shrinking lipid vesicle. Phys Rev E. 2009;80(5).

14. Uchino T, Matsumoto Y, Murata A, Oka T, Miyazaki Y, Kagawa Y. Transdermal delivery of flurbiprofen from surfactant-based vesicles: Particle characterization and the effect of water on in vitro transport. Int J Pharmaceut. 2014;464(1-2):75–84.

15. Ge XM, Wei MY, He SN, Yuan WE. Advances of Non-Ionic Surfactant Vesicles (Niosomes) and Their Application in Drug Delivery. Pharmaceutics. 2019;11(2).

16. Ilhan-Ayisigi E, Yaldiz B, Bor G, Yaghmur A, Yesil-Celiktas O. Advances in microfluidic synthesis and coupling with synchrotron SAXS for continuous production and real-time structural characterization of nano-self-assemblies. Colloid Surface B. 2021;201.

17. Elsayed MMA, Ibrahim MM, Cevc G. The effect of membrane softeners on rigidity of lipid vesicle bilayers: Derivation from vesicle size changes. Chem Phys Lipids. 2018;210:98–108.

18. Riske KA, Domingues CC, Casadei BR, Mattei B, Carita AC, Lira RB, et al. Biophysical approaches in the study of biomembrane solubilization: quantitative assessment and the role of lateral inhomogeneity. Biophys Rev. 2017;9(5):649–67.

19. Seddon AM, Curnow P, Booth PJ. Membrane proteins, lipids and detergents: not just a soap opera. Biochim Biophys Acta. 2004;1666(1-2):105–17.

20. Lichtenberg D, Ahyayauch H, Goni FM. The mechanism of detergent solubilization of lipid bilayers. Biophys J. 2013;105(2):289–99.

21. Helenius A, Simons K. Solubilization of membranes by detergents. Biochim Biophys Acta. 1975;415(1):29–79.

22. Sujatha J, Mishra AK. Effect of ionic and neutral surfactants on the properties of phospholipid vesicles: Investigation using fluorescent probes. J Photoch Photobio A. 1997;104(1-3):173–8.

23. Dalgarno PA, Juan-Colas J, Hedley GJ, Pineiro L, Novo M, Perez-Gonzalez C, et al. Unveiling the multi-step solubilization mechanism of sub-micron size vesicles by detergents. Sci Rep-Uk. 2019;9.

24. Juan-Colas J, Dresser L, Morris K, Lagadou H, Ward RH, Burns A, et al. The Mechanism of Vesicle Solubilization by the Detergent Sodium Dodecyl Sulfate. Langmuir. 2020;36(39):11499–507.

25. Antonny B. Mechanisms of Membrane Curvature Sensing. Annu Rev Biochem. 2011;80:101–23.

26. Shibata Y, Hu JJ, Kozlov MM, Rapoport TA. Mechanisms Shaping the Membranes of Cellular Organelles. Annu Rev Cell Dev Bi. 2009;25:329–54.

27. Niroomand H, Venkatesan GA, Sarles SA, Mukherjee D, Khomami B. Lipid-Detergent Phase Transitions During Detergent-Mediated Liposome Solubilization. J Membrane Biol. 2016;249(4):523–38.

28. Goni FM, Alonso A. Spectroscopic techniques in the study of membrane solubilization, reconstitution and permeabilization by detergents. Bba-Biomembranes. 2000;1508(1-2):51–68.

29. Muddana HS, Chiang HH, Butler PJ. Tuning membrane phase separation using nonlipid amphiphiles (vol 102, pg 489, 2012). Biophysical Journal. 2012;103(4):846-.

30. Pizzirusso A, De Nicola A, Milano G. MARTINI Coarse-Grained Model of Triton TX-100 in Pure DPPC Monolayer and Bilayer Interfaces. J Phys Chem B. 2016;120(16):3821–32.

31. Bandyopadhyay S, Shelley JC, Klein ML. Molecular dynamics study of the effect of surfactant on a biomembrane. J Phys Chem B. 2001;105(25):5979–86.

32. Kamrath RF, Frances EI. Mass-action model of mixed micellization. J Phys Chem. 1984;88(8):1642–8.

33. Flagmeier P, De S, Wirthensohn DC, Lee SF, Vincke C, Muyldermans S, et al. Ultrasensitive measurement of Ca2+ influx into lipid vesicles induced by protein aggregates. Eur Biophys J Biophy. 2017;46:S237–S.

34. Knoch H, Ulbrich MH, Mittag JJ, Buske J, Garidel P, Heerklotz H. Complex Micellization Behavior of the Polysorbates Tween-20 and Tween 80. Mol Pharm. 2021;18(8):3147–57.

35. Gupta A, Korte T, Herrmann A, Wohland T. Plasma membrane asymmetry of lipid organization: fluorescence lifetime microscopy and correlation spectroscopy analysis. J Lipid Res. 2020;61(2):252–66.

36. Stockl M, Plazzo AP, Korte T, Herrmann A. Detection of lipid domains in model and cell membranes by fluorescence lifetime imaging microscopy of fluorescent lipid analogues. J Biol Chem. 2008;283(45):30828–37.

37. Schroter F, Jakop U, Teichmann A, Haralampiev I, Tannert A, Wiesner B, et al. Lipid dynamics in boar sperm studied by advanced fluorescence imaging techniques. Eur Biophys J Biophy. 2016;45(2):149–63.

38. Bag N, Yap DHX, Wohland T. Temperature dependence of diffusion in model and live cell membranes characterized by imaging fluorescence correlation spectroscopy. Bba-Biomembranes. 2014;1838(3):802–13.

39. Flagmeier P, De SM, Michaels TCT, Yang XT, Dear AJ, Emanuelsson C, et al. Direct measurement of lipid membrane disruption connects kinetics and toxicity of A beta 42 aggregation. Nat Struct Mol Biol. 2020;27(10):886-+.

40. Palmieri V, Lucchetti D, Gatto I, Maiorana A, Marcantoni M, Maulucci G, et al. Dynamic light scattering for the characterization and counting of extracellular vesicles: a powerful noninvasive tool. J Nanopart Res. 2014;16(9).

41. Sarra A, Celluzzi A, Bruno SP, Ricci C, Sennato S, Ortore MG, et al. Biophysical Characterization of Membrane Phase Transition Profiles for the Discrimination of Outer Membrane Vesicles (OMVs) From Escherichia coli Grown at Different Temperatures. Front Microbiol. 2020;11.

42. Dresser L, Hunter P, Yendybayeva F, Hargreaves AL, Howard JAL, Evans GJO, et al. Amyloid-beta oligomerization monitored by single-molecule stepwise photobleaching. Methods. 2021;193:80–95.

43. Shi X, Lim J, Ha T. Acidification of the oxygen scavenging system in single-molecule fluorescence studies: in situ sensing with a ratiometric dual-emission probe. Anal Chem. 2010;82(14):6132–8.

44. Roy R, Hohng S, Ha T. A practical guide to single-molecule FRET. Nat Methods. 2008;5(6):507–16.

45. Cimorelli M, Nieuwland R, Varga Z, van der Pol E. Standardized procedure to measure the size distribution of extracellular vesicles together with other particles in biofluids with microfluidic resistive pulse sensing. Plos One. 2021;16(4).

46. Pfrieger FW, Vitale N. Cholesterol and the journey of extracellular vesicles. J Lipid Res. 2018;59(12):2255–61.

47. Conteduca D, Quinn SD, Krauss TF. Dielectric metasurface for high-precision detection of large unilamellar vesicles. J Optics-Uk. 2021;23(11).

48. Chuo ST, Chien JC, Lai CP. Imaging extracellular vesicles: current and emerging methods. J Biomed Sci. 2018;25(1):91.

49. Wu YT, Deng WT, Klinke DJ. Exosomes: improved methods to characterize their morphology, RNA content, and surface protein biomarkers. Analyst. 2015;140(19):6631–42.

50. Nguyen DB, Ly TBT, Wesseling MC, Hittinger M, Torge A, Devitt A, et al. Characterization of Microvesicles Released from Human Red Blood Cells. Cell Physiol Biochem. 2016;38(3):1085–99.

51. Balomenos AD, Stefanou V, Manolakos ES. Analytics and visualization tools to characterize single-cell stochasticity using bacterial single-cell movie cytometry data. Bmc Bioinformatics. 2021;22(1).

52. Karatekin E, Sandre O, Guitouni H, Borghi N, Puech PH, Brochard-Wyart F. Cascades of transient pores in giant vesicles: Line tension and transport. Biophysical Journal. 2003;84(3):1734–49.

53. Colom A, Derivery E, Soleimanpour S, Tomba C, Molin MD, Sakai N, et al. A fluorescent membrane tension probe. Nat Chem. 2018;10(11):1118–25.

54. Osteikoetxea X, Sodar B, Nemeth A, Szabo-Taylor K, Paloczi K, Vukman KV, et al. Differential detergent sensitivity of extracellular vesicle subpopulations. Org Biomol Chem. 2015;13(38):9775–82.

55. Hannestad JK, Rocha S, Agnarsson B, Zhdanov VP, Wittung-Stafshede P, Hook F. Single-vesicle imaging reveals lipid-selective and stepwise membrane disruption by monomeric alpha-synuclein. Proc Natl Acad Sci U S A. 2020;117(25):14178–86.

56. Armstrong JK, Wenby RB, Meiselman HJ, Fisher TC. Vybrant (TM) DiO, DiI and DiD dyes for multiple labeling of red blood cell populations for in vivo survival studies. Blood. 2004;104(11):442a–a.

57. Zhang Z, Yomo D, Gradinaru C. Choosing the right fluorophore for single-molecule fluorescence studies in a lipid environment. Biochim Biophys Acta Biomembr. 2017;1859(7):1242–53.

58. De S, Wirthensohn DC, Flagmeier P, Hughes C, Aprile FA, Ruggeri FS, et al. Different soluble aggregates of Abeta42 can give rise to cellular toxicity through different mechanisms. Nat Commun. 2019;10(1):1541.

59. Miller H, Cosgrove J, Wollman AJM, Taylor E, Zhou Z, O’Toole PJ, et al. High-Speed Single-Molecule Tracking of CXCL13 in the B-Follicle. Front Immunol. 2018;9:1073.

60. Preus S, Noer SL, Hildebrandt LL, Gudnason D, Birkedal V. iSMS: single-molecule FRET microscopy software. Nat Methods. 2015;12(7):593–4.

