## Supplementary for "Tween-20 induces the structural remodelling of single lipid vesicles"

### SUPPORTING INFORMATION

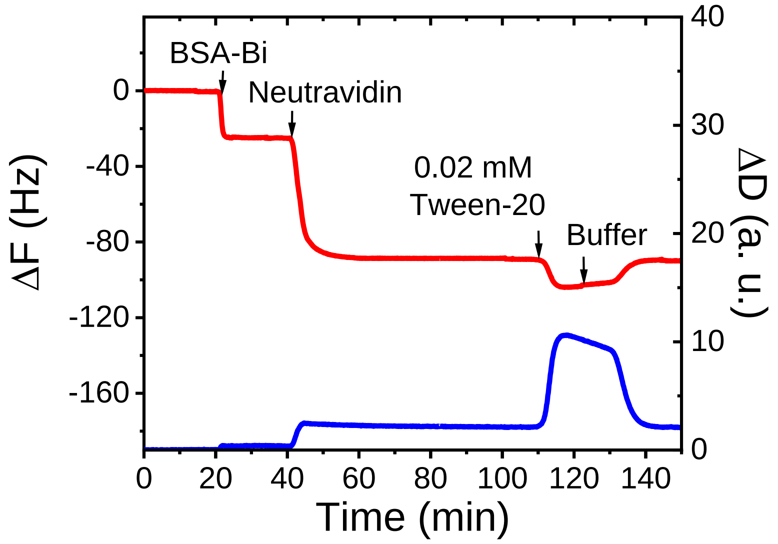


**Figure S1. The non-specific and reversible attachment of Tween-20 to a SiO2 sensor surface containing BSA, BSA-Biotin and Neutravidin.** The time evolution of ΔF and ΔD upon the addition of BSA-Biotin, Neutravidin and 0.02 mM Tween-20 to a $SiO_2$ sensor surface at 21$^{o}$C. After saturation of the surface by Tween-20 was reached at t = 120 min, the sensor was flushed with 50 mM Tris buffer (pH 8).

#### Supplementary Text 1: Mass-Action Model for Mixed Micellization.

The ensemble FRET efficiency of POPC vesicles is interpreted to be proportional to the surface concentration of interacting DiI and DiD in the membrane, [ DiI*DiD ]=K[ DiI ][ DiD ] relative to the maximum surface concentration of interaction pairs, min( [ DiI ], [ DiD ] ). In our experiments both concentrations are equal, [ Di ]=[ DiI ]=[ DiD ], so $E_{FRET} \propto [Di]$. A decrease in the FRET efficiency occurs if this surface concentration decreases, which occurs upon swelling of the vesicles when Tween-20 molecules are incorporated in the membrane. Hence, we may write

$$E_{FRET} = A + B\phi, \qquad \text{(Equation 1)}$$

where *A* and *B* are constants, $\phi$ is the volume fraction of lipids, and 1-$\phi$ is the volume fraction of Tween-20 in the membrane (assuming the concentration of FRET tags is very low). In this section we focus on the molecular mechanism by which $\phi$ depends on the experimentally tunable overall concentration of Tween-20.

In previous work[1], we heuristically applied the Hill model to describe the self-assembly curves, $\phi$=1/(1+([Tween-20]/k)$^n$), where *k* is the half maximal concentration and *n* is the Hill coefficient, which controls how sharp the self-assembly transition is at high concentrations. This approach has been useful to characterise a solubilization concentration (in terms of the half maximal concentration) and the sharpness of the transition, but gives a poor fit quality at low concentrations below the transition, which indicates there are some pieces of physics missing. Indeed, the Hill model assumes that groups of *n* molecules cooperatively adsorb to a limited number of binding sites at the vesicle without taking into account that 1. the molecules may adsorb independently, 2. the adsorbed molecules have some translational entropy in the membrane and exhibit interactions with other adsorbed molecules and the lipids, 3. the membrane does not have a limited number of binding sites and may in principle swell indefinitely. Hence, in the present work we develop a simple mass-action model for mixed micellization that takes this physics into account[2].

As a starting point, we consider a solution in a volume *V* at temperature *T* with a total number density of lipids, $\rho_L$ (we presume all lipids to reside in the membrane), and a total number density Tween-20, $\rho_T = N_A[Tween-20]$(with $N_A$ Avogadro's constant), which is partitioned into a concentration that is free in solution, $\rho_{T,s}$, and a concentration that is adsorbed to the vesicle $\rho_{T,v}$, hence, $\rho_T = \rho_{T,s} + \rho_{T,v}$, and the volume fraction of lipids in the membrane is $\phi = \rho_{L,v}/(\rho_{T,v} + \rho_L)$. We will now formulate the free-energy contributions of the lipids and of freely dissolved and adsorbed Tween-20, and minimise the overall free energy to obtain the dependence of $\phi$ on $\rho_T$via $\rho_{T,v}$.

A Tween-20 molecule that is free in solution has translational entropy, which provides a free energy contribution $kTln(\rho_{T,v}V_m) - kT$, with *k* Boltzmann's constant and with $V_m$ the molecular volume. To adsorb to the vesicle, it loses this translational entropy, but gains a binding enthalpy *h*<0, as well as translational entropy within in the membrane. If self-interactions are preferred over interactions with the lipids, this repulsive interaction is captured using the mean-field interaction energy $\phi(1-\phi)\chi kT$ per molecule, with $\chi$ the Flory-Huggins parameter. This free energy of mixing,

$$f_{mix}(\phi) = [\phi ln\phi + (1-\phi)ln(1-\phi)]kT + \phi(1-\phi)\chi kT \qquad \text{(Equation 2)}$$

is contributed to by both the Tween-20 molecules and the lipids in the membrane. Finally, we take into account that each Tween-20 molecule has a chemical potential *μ*, which acts as a Lagrange multiplier, as it enables us to minimise the total free energy density

$$\frac{F}{V} = \rho_{T,s}\left[kTln\left(\rho_{T,s}V_m\right) - kT - \mu\right] + \rho_{T,v}\left[f_{mix}(\phi) + h - \mu\right] + \rho_L f_{mix}(\phi) \qquad \text{(Equation 3)}$$

with respect to $\rho_{T,s}$ and $\rho_{T,v}$independently, and the constraint $\rho = \rho_{T,s} + \rho_{T,v}$ is obeyed by adjusting $\mu$ accordingly. Indeed, by minimising the free energy density with respect to $\rho_{T,s}$ we obtain $\mu = kTln\left(\rho_{T,s}V_m\right)$, and minimization with respect to $\rho_{T,v}$ gives $0 = f_{mix} + h + \left(\rho_{T,v} + \rho_L\right)\frac{df_{mix}}{d\rho_{T,v}} - \mu$ . After substituting the chemical potential, and using the product rule, $df_{mix}/d\rho_{T,v} = (df_{mix}/d\phi)\left(d\,\phi/d\,\rho_{T,v}\right)$, this gives $0 = ln(1-\phi) + \phi^2\chi + h/kT - ln\left(\rho_{T,s}V_m\right)$ . We now introduce the equilibrium constant for binding, $K=V_m \exp(-h/kT) \equiv V_m \exp(-\Delta H/RT)$, with $\Delta H$ the binding enthalpy and $R=kN_A$ the gas constant, and substitute $\rho_{T,s} = \rho_T - \rho_L\,(1-\phi)/\phi$, to obtain the final result

$$Kexp(-\phi^2\chi)\left(\frac{\rho_T}{1-\phi} - \frac{\rho_L}{\phi}\right) = 1 \qquad \text{(Equation 4)}$$

which is an implicit equation for $\phi$ as a function of $\rho_T = N_A[Tween-20]$, with as parameters the equilibrium constant $K$, interaction parameter $\chi$, and the overall lipid concentration $\rho_L$.

To inspect how these parameters affect the titration curve, we first focus on the asymptotic limit where Tween-20 and the lipid are perfectly miscible in the membrane, i.e., we consider the case $\chi=0$. Here, the volume fraction of lipids in the membrane is given by

$$\phi = \frac{1}{2}\left(1 - K(\rho_T + \rho_L) + \sqrt{(1 - K(\rho_T + \rho_L)^2 + 4K\rho_L)}\right) \qquad \text{(Equation 5)}$$

In **Figure S2** we plot this curve for various lipid concentrations $\rho_L$. We find that this concentration shifts the half-maximal concentration as $\rho_{1/2} = \rho_L + 1/(2K)$ (Hence, if the data would be analysed using the Hill model, this would yield $k \approx (1 + 2\rho_L K)/2\,K$). Further, we find that the transition is broad and smooth at high lipid concentrations, but that the transition becomes sharp at $K\rho_T = 1$ for vanishing lipid concentrations. Within the Hill model, this behaviour would be attributed to an increasing cooperativity of Tween-20 adsorption; our new interpretation is that a sharper transition indicates a larger excess of Tween-20 relative to the number of lipid molecules at the transition.

The influence of the interaction parameter can be assessed in the regime where it remains small, $\chi$<<1. Here, $exp(-(1-\phi)^2\chi) \approx -(1-\phi)^2\chi$ can be inserted, which yields the cubic equation $K(\rho_T + \rho_L)\phi^3 - (1 + K\rho_L)\phi^2 + \left(1 - K(\rho_T + \rho_L)\right)\phi + K\rho_L = 0$ . For low lipid concentrations, again the transition becomes sharp ($\phi$=0 for $K\rho_T > 1$), and below the transition concentration, $K\rho_T < 1$, we have

$$\phi = \frac{1}{2\chi K\rho_T}\left(1 - \sqrt{1 - 4\chi K\rho_T(1 - K\rho_T)}\right), \qquad \text{(Equation 6)}$$

which we plot in **Figure S2**. For low concentrations, $K\rho_T \ll 1$, this gives $\phi = 1 - (1-\chi)K\rho_T$, and resembles the non-cooperative Hill model (i.e., for $n$=1) if $\chi$=0. Hence, from the shape of the curve we assess information about the interaction of Tween-20 with the lipids.

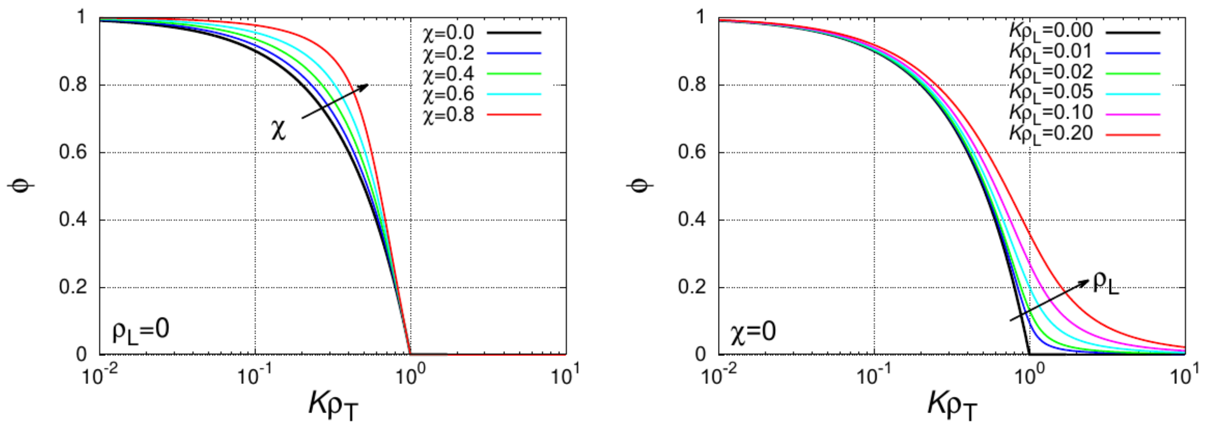


**Figure S2. Adsorption isotherm of Tween-20 to a vesicle.** The volume fraction of lipids in a membrane, $\phi$, against the Tween-20 concentration, $\rho_T$. The curves are controlled by the equilibrium constant $K$, the concentration of lipids, $\rho_L$, and by the interaction parameter $\chi$. For low lipid concentrations, $\rho_L \rightarrow 0$, there is a sharp transition, and the shape of the curve is controlled by $\chi$. For ideal mixing of Tween-20 with the lipids, $\chi$=0, the transition becomes less sharp with an increasing lipid concentration.

We fit both the Hill model and the mass-action model to our $E_{FRET}$ data to assess the fit quality and extract the physical parameter values. First, we determine the baseline value $A$ in Eq. (1) at high concentrations. From this we obtain the experimental variance $\sigma_E^2$. We then use this to calculate the reduced chi square definition

$$\chi^2 = \frac{1}{N_{data} - N_{par}} \sum_{i=1}^{N_{data}} \frac{\left(E_{i,fit} - E_{i,data}\right)^2}{\sigma_E^2} \qquad \text{(Equation 7)}$$

where $N_{par}$=3 for the Hill model (which has parameters $B$, $k$, and $n$) and $N_{par}$=4 for the mass-action model (which has parameters $B$, $K$, $\chi$, $\rho_L$). We note that $\rho_L$ is the concentration of lipids, which is strictly known.


1. Dalgarno PA, Juan-Colas J, Hedley GJ, Pineiro L, Novo M, Perez-Gonzalez C, et al. Unveiling the multi-step solubilization mechanism of sub-micron size vesicles by detergents. Sci Rep-Uk. 2019;9.
2. Kamrath RF, Franses EI. Mass-Action Model of Mixed Micellization, J. Phys. Chem. 1985;88:1642

**Table S1.** Fitting parameters associated with mass action model fits applied to the $E_{FRET}$ titrations shown in Figure 2(B). Using the Van 't Hoff relationship $K \propto exp(\Delta H/RT)$for the equilibrium constant K, we found an enthalpy $\Delta H$=– 31 +/- 3 kJ/mol for binding surfactants to a lipid membrane.

| | **4 °C** | **21 °C** | **37 °C** |
|---|---|---|---|
| A | 0.055 ± 0.002 | 0.55 ± 0.002 | 0.55 ± 0.002 |
| B | 0.382 ± 0.007 | 0.382 ± 0.007 | 0.382 ± 0.007 |
| $\rho_L$ (mM) | 0.0022 ± 0.0004 | 0.0022 ± 0.0004 | 0.0022 ± 0.0004 |
| $K$ ($mM^{-1}$) | 6.27 ± 0.11 | 11.93 ± 0.32 | 26.39 ± 1.06 |
| $\chi$ | 1.15 ± 0.12 | 1.22 ± 0.17 | 1.24 ± 0.21 |

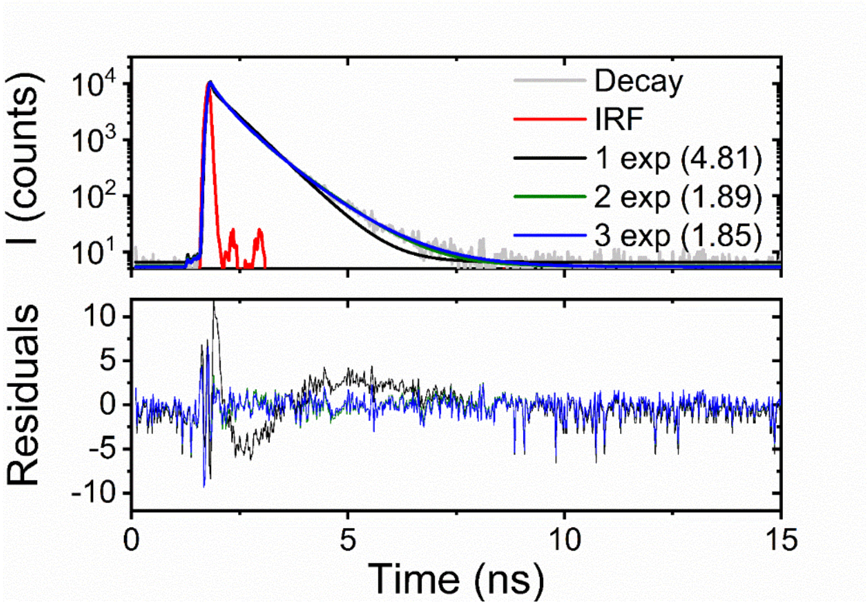


**Figure S3. Time-resolved fluorescence decays obtained from DiI and DiD loaded vesicles fit to a tri-exponential model.** Top panel: Representative time-resolved fluorescence decay (grey) obtained from DiI and DiD coated POPC vesicles in the absence of Tween-20. Black, green and blue solid lines represent reconvolution fits to mono-, bi- and tri-exponential decay functions, respectively. The red solid line represents the instrument response function (IRF). $\lambda_{ex}$ = 532 nm (50 MHz). The numbers in brackets represent goodness of fit, $\chi^2$, values. Bottom panel: residuals obtained for the mono-, bi- and tri-exponential decay functions. Solution conditions:25 μM POPC, 0.025 μM DiI, 0.025 μM DiD, 50 mM Tris, pH 8.0.

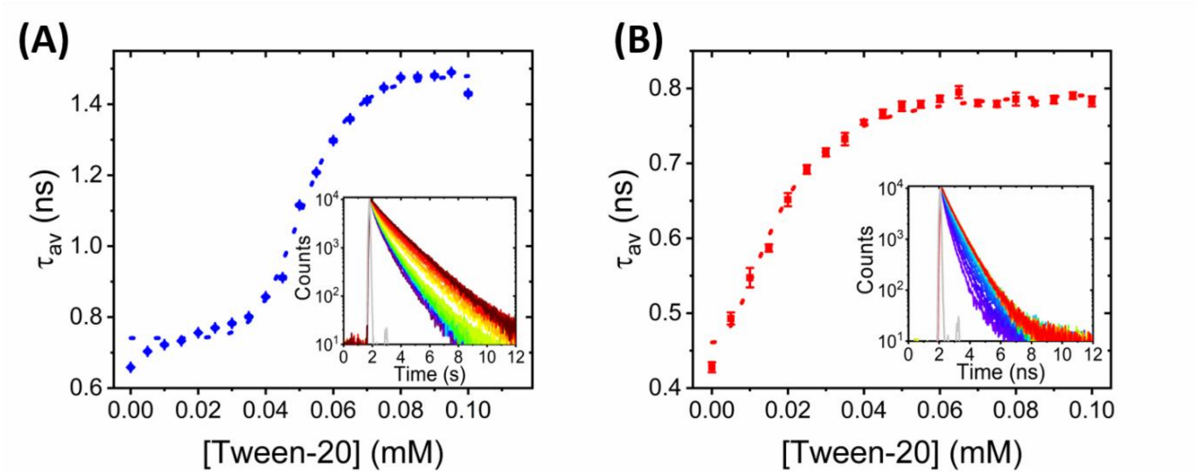


**Figure S4. Time-resolved fluorescence decays obtained from DiI and DiD loaded vesicles as a function of Tween-20 at 4ºC and 37ºC.** The amplitude weighted average lifetime of DiI in the presence of DiD as a function of Tween-20 at (A) 4ºC and (B) 37ºC with $\lambda_{ex}$ = 532 nm (50 MHz). Insets: the corresponding time-resolved fluorescence decays. The instrumental response function acquired under both conditions is shown in grey. Solution conditions: 25 μM POPC, 0.025 μM DiI, 0.025 μM DiD, 50 mM Tris, pH 8.0.

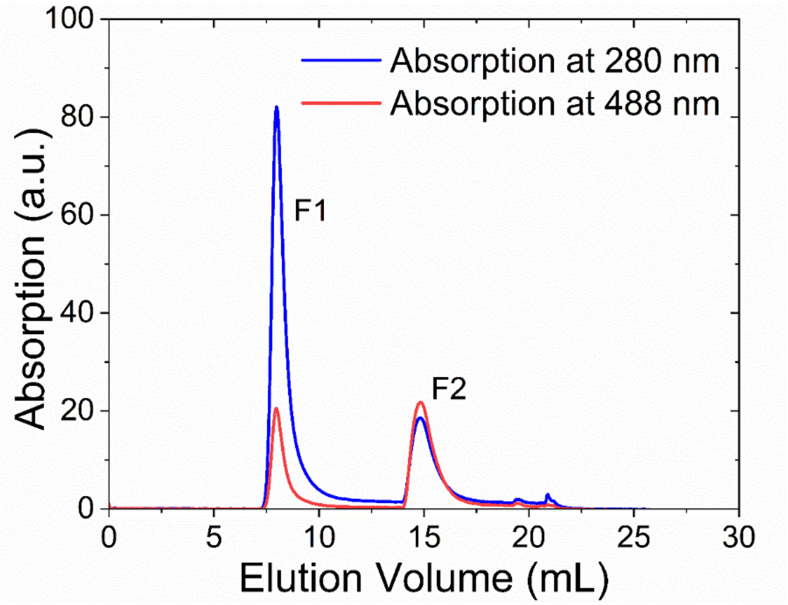


**Figure S5. Separation of Cal-520 loaded POPC vesicles from non-incorporated Cal-520 via size exclusion chromatography.** 200 nm sized POPC vesicles were prepared in 50 mM Tris buffer (pH 8.0) with 100 mM Cal-520 by the extrusion method. Size-exclusion chromatography was then performed using a 10/30 column attached to an AKTA pure system (GE Healthcare). Absorption was monitored at 280 nm (blue) and 488 nm (red). Fractions F1 and F2 represent the vesicle fraction and free dye/lipid fraction respectively.

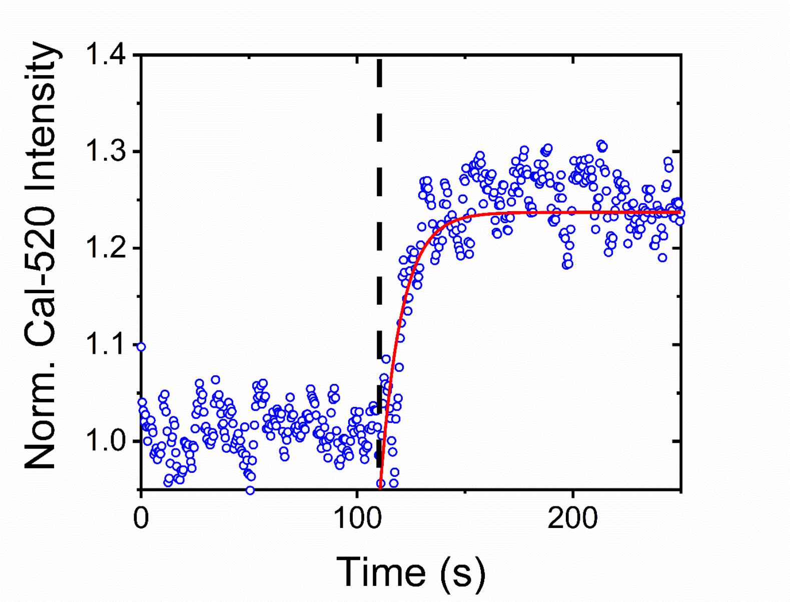


**Figure S6. Kinetics of Tween-20 induced membrane disruption followed by the fluorescent enhancement of Cal-520.** Normalized variation in the fluorescence emission intensity obtained from POPC vesicles encapsulating Cal-520 before (< 120 s) and after (> 120 s) injection of 0.2 mM Tween-20. The dashed line represents the point of Tween-20 injection and the solid red line represents a mono-exponential fit.

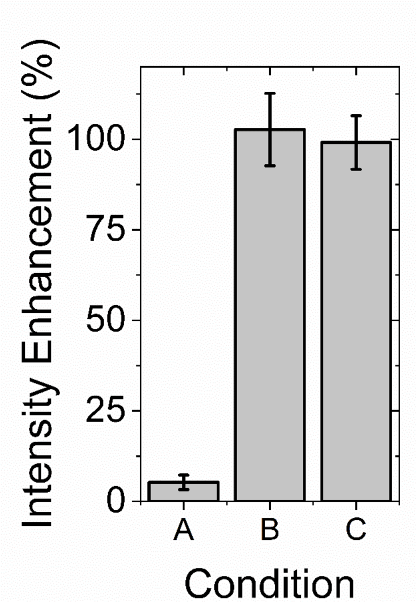


**Figure S7.** Comparative bar plot summarizing the relative variation in fluorescence intensity enhancement observed from Cal-520 loaded POPC vesicles in (A) 50 mM Tris (pH 8), 1 mM $Ca^{2+}$, (B) 50 mM Tris (pH 8), 1 mM $Ca^{2+}$, 1 mg/mL ionomycin and (C) 50 mM Tris (pH 8), 1 mM $Ca^{2+}$, 0.06 mM Tween-20. Error bars represent the standard error of the mean from 3 experimental runs. Solution conditions: 25 μM POPC, 1 μM Cal-520.

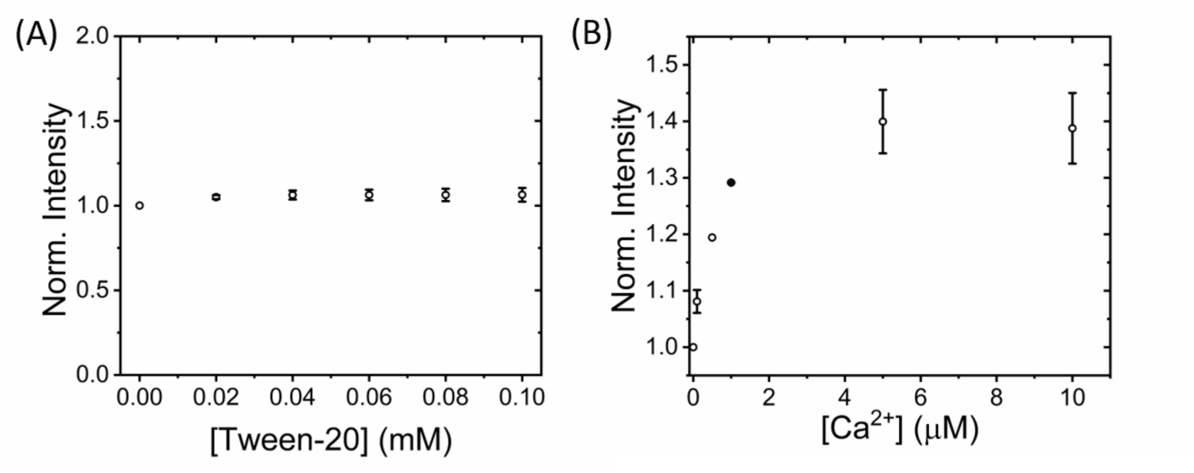


**Figure S8. Representative variation in free Cal-520 emission intensity.** Normalized variation in the fluorescence emission intensity of 0.1 μM Cal-520 as (A) a function of Tween-20 and (B) as a function of $Ca^{2+}$ in the presence of 0.2 mM Tween-20. In both cases, the base buffer was 50 mM Tris, pH 8.0, 21°C.

**Table S2.** Fitting parameters and errors associated with Hill Model fits applied to the Cal-520-loaded vesicle titrations shown in Figure 2(D).

| | **4 °C** | **21 °C** | **37 °C** |
|---|---|---|---|
| A | 1.04 ± 0.02 | 1.04 ± 0.06 | 1.03 ± 0.08 |
| B | 1.92 ± 0.03 | 1.84 ± 0.02 | 1.93 ± 0.04 |
| k (mM) | 0.05 ± 0.01 | 0.015 ± 0.01 | 0.022 ± 0.003 |
| N | 2.75 ± 0.28 | 2.53 ± 0.43 | 1.90 ± 0.39 |
| $R^2$ | 0.99 | 0.98 | 0.97 |

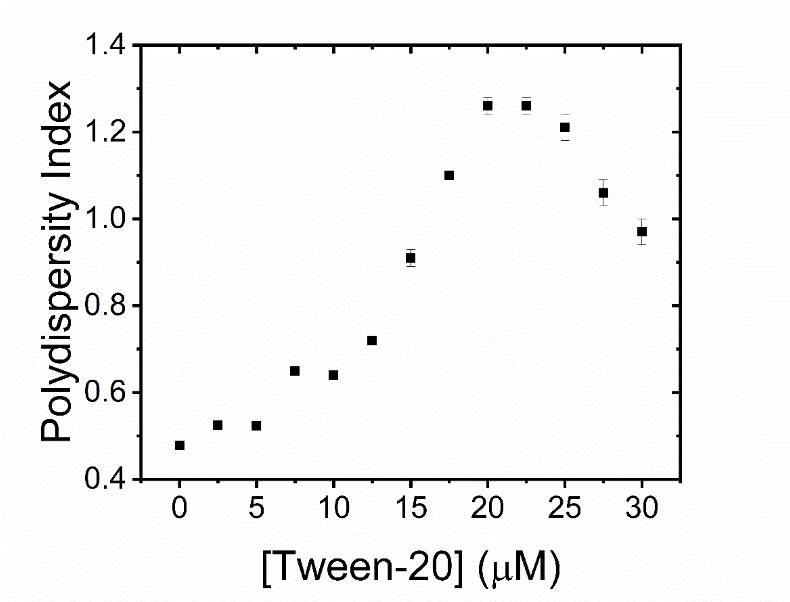


**Figure S9. Polydispersity index of POPC vesicles in the presence of Tween-20.** Variation in the polydispersity index of POPC vesicles suspended in 50 mM Tris (pH 8.0) buffer in the absence and presence of Tween-20. Data points represent the mean (± SEM) from 3 separate experimental runs. Solution conditions: 25 μM POPC, 0.025 μM DiI, 0.025 μM DiD, 50 mM Tris, pH 8.0.

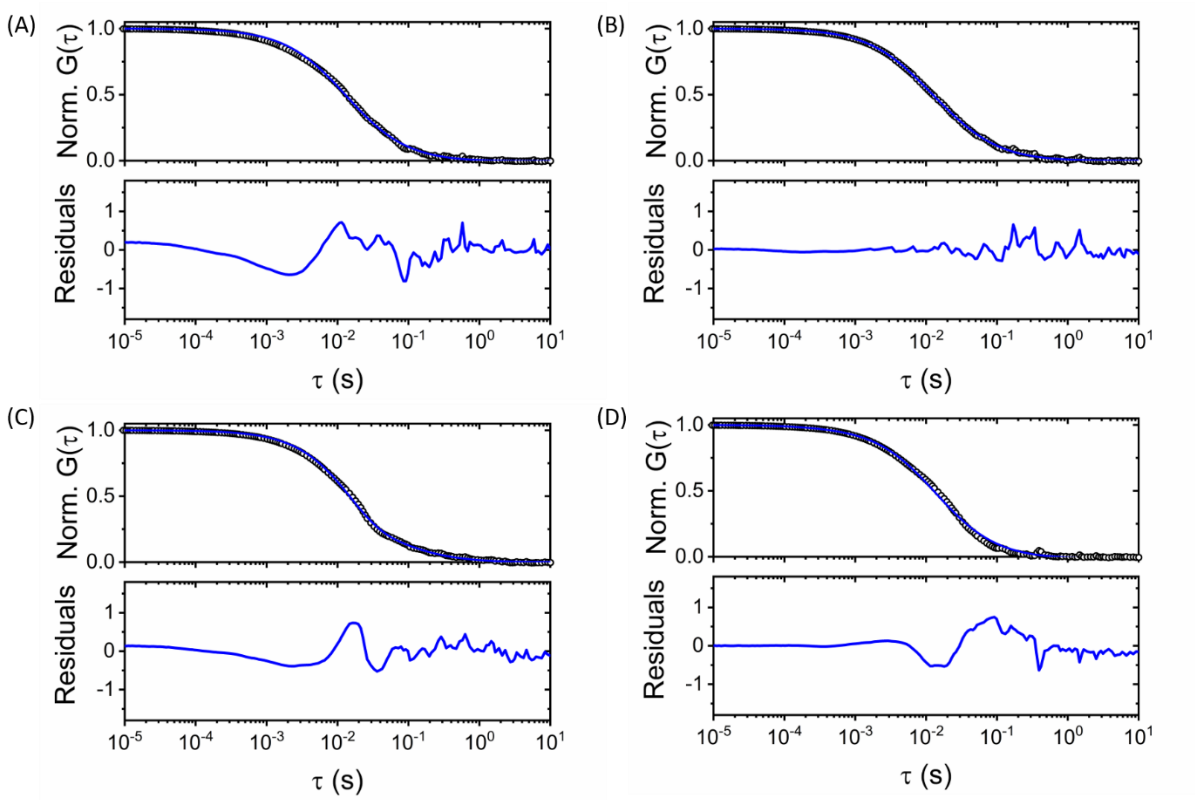


**Figure S10. Tween-20 vesicle interactions reported using FCS.** Cross correlation curves (black) and fits (blue) (top panel) associated with 200 nm sized POPC vesicles containing 0.1 % DiI in the presence of (A) 0 mM, (B) 0.02 mM, (C) 0.04 mM and (D) 0.06 mM Tween-20. Bottom panels represent residuals of the fits.

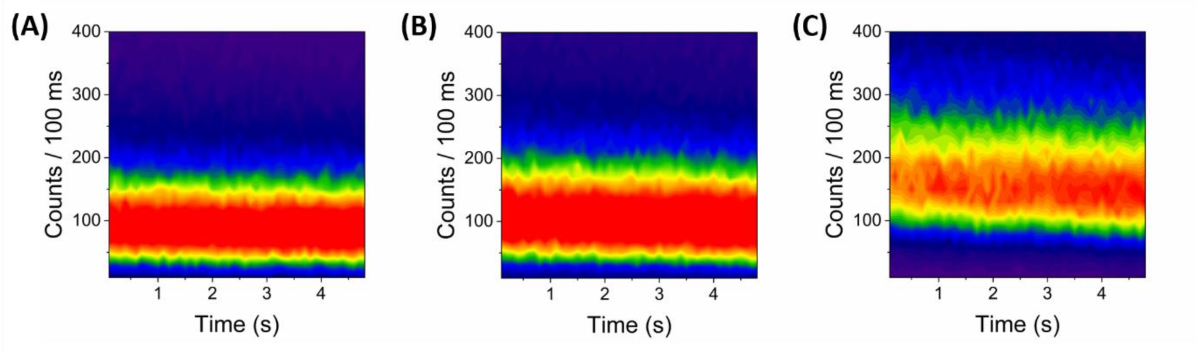


**Figure S11. Photostability of Cal-520 loaded vesicles in the absence and presence of Tween-20.** Contour plots of the time-evolution of Cal-520 vesicle population in (A) 50 mM Tris buffer, pH 8, (B) 50 mM Tris buffer, 10 mM $Ca^{2+}$, pH 8 and (C) 50 mM Tris buffer, 10 mM $Ca^{2+}$, 0.01 mM Tween-20, pH 8. Each contour plot was produced by superimposing individual intensity trajectories and contours are plotted from blue (lowest population) to red (highest population).

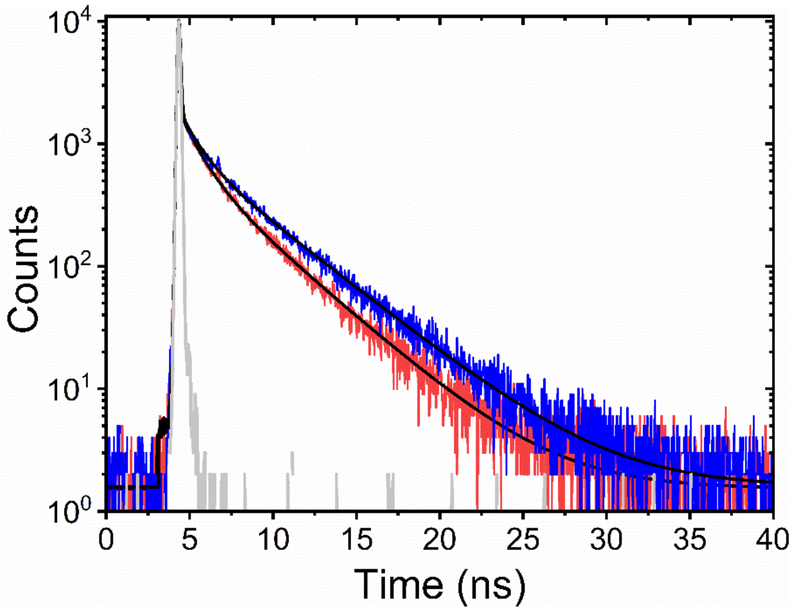


**Figure S12. Reduction in membrane tension monitored by FliptR response.** Time-resolved fluorescence decays obtained from POPC vesicles coated in FliptR in the absence (blue) and presence of 0.5 mM Tween-20 (red). Solid black lines represent tri-exponential fits to the experimental data. Also shown is the instrumental response function (grey). Solution conditions: 25 μM POPC, 0.4 μM FliptR, 50 mM Tris, pH 8.
